# Pharmacological targeting of a PWWP domain demonstrates cooperative control of NSD2 localization

**DOI:** 10.1101/2021.03.05.433782

**Authors:** David Dilworth, Ronan P. Hanley, Renato Ferreira de Freitas, Abdellah Allali-Hassani, Mengqi Zhou, Naimee Mehta, Matthew R. Marunde, Suzanne Ackloo, Edyta Marcon, Fengling Li, Irene Chau, Albina Bolotokova, Su Qin, Ming Lei, Yanli Liu, Magdalena M Szewczyk, Aiping Dong, Sina Kazemzadeh, Tigran Abramyan, Irina K Popova, Nathan W Hall, Matthew J Meiners, Marcus A Cheek, Elisa Gibson, Dmitri Kireev, Jack F. Greenblatt, Michael-C. Keogh, Jinrong Min, Peter J. Brown, Masoud Vedadi, Cheryl H. Arrowsmith, Dalia Barsyte-Lovejoy, Lindsey I James, Matthieu Schapira

## Abstract

NSD2 is the primary enzyme responsible for the dimethylation of lysine 36 of histone 3 (H3K36me2), a mark associated with active gene transcription and intergenic DNA methylation. In addition to a methyltransferase domain, NSD2 harbors two PWWP and five PHD domains believed to serve as chromatin reading modules, but their exact function in the regulation of NSD2 activity remains underexplored. Here we report a first-in-class chemical probe targeting the N-terminal PWWP (PWWP1) domain of NSD2. **UNC6934** binds potently (K_d_ of 91 ± 8 nM) to PWWP1, antagonizes its interaction with nucleosomal H3K36me2, and selectively engages endogenous NSD2 in cells. Crystal structures show that **UNC6934** occupies the canonical H3K36me2-binding pocket of PWWP1 which is juxtaposed to the DNA-binding surface. In cells, **UNC6934** induces accumulation of endogenous NSD2 in the nucleolus, phenocopying the localization defects of NSD2 protein isoforms lacking PWWP1 as a result of translocations prevalent in multiple myeloma. Mutation of other NSD2 chromatin reader domains also increases NSD2 nucleolar localization, and enhances the effect of **UNC6934**. Finally we identified two C-terminal nucleolar localization sequences in NSD2 that appear to drive nucleolar accumulation when one or more chromatin reader domains are disabled. These data support a model in which NSD2 chromatin engagement is achieved in a cooperative manner and subcellular localization is controlled by multiple competitive structural determinants. This chemical probe and the accompanying negative control, **UNC7145**, will be useful tools in defining NSD2 biology.

## Introduction

Nuclear receptor-binding SET domain-containing 2 (NSD2, also known as MMSET and WHSC1) is a protein lysine methyltransferase that belongs to the NSD family, which also includes NSD1 and NSD3. NSD2 predominantly dimethylates lysine 36 of histone 3 (H3K36)^1^ and is aberrantly expressed, amplified or somatically mutated in multiple types of cancer^2^. Notably, the t(4;14) NSD2 translocation in multiple myeloma (MM) and the hyper-activating NSD2 E1099K mutation in a subset of pediatric acute lymphoblastic leukemia (ALL) both result in altered chromatin methylation which drives oncogenesis^1,3-7^. Functionally, NSD2 is responsible for the bulk of H3K36me2 in diverse cell types. Dimethylation of H3K36 by both NSD1 and NSD2 recruits DNMT3a at intergenic regions to control DNA methylation and regulate development and homeostasis^8,9^. NSD2 is also required for efficient non-homologous end-joining and homologous recombination, two canonical DNA repair pathways^10,11^.

In addition to its catalytic domain, NSD2 has multiple protein-protein interaction (PPI) modules with known or potential chromatin reading functions, including five PHD (plant homeodomain) and two PWWP (proline-tryptophan-tryptophan-proline) domains^2^, as well as a putative DNA-binding HMG-box (high mobility group box) domain (**Fig. 1a**). Mounting evidence suggests that these domains play important roles in NSD2 function, but the individual and/or collective roles of the NSD2 chromatin reader domains are still being elucidated^2^. Many PWWP domains are known H3K36me2,3 reading modules that engage methyl-lysine while simultaneously interacting with nucleosomal DNA adjacent to H3K36^12,13^. The isolated N-terminal PWWP domain of NSD2 (NSD2-PWWP1) binds H3K36 di- and trimethylated nucleosomes; this interaction presumably is mediated by a conserved aromatic cage and stabilizes NSD2 at chromatin^8^. Mutation of the aromatic cage residues abrogates NSD2-PWWP1 binding to nucleosomal H3K36me2, but has only modest effect on global H3K36 methylation level in cells^8^. However, H3K36 methylation has been shown to be abolished upon mutation of the second PHD domain (PHD2)^14^.

**Figure 1:**
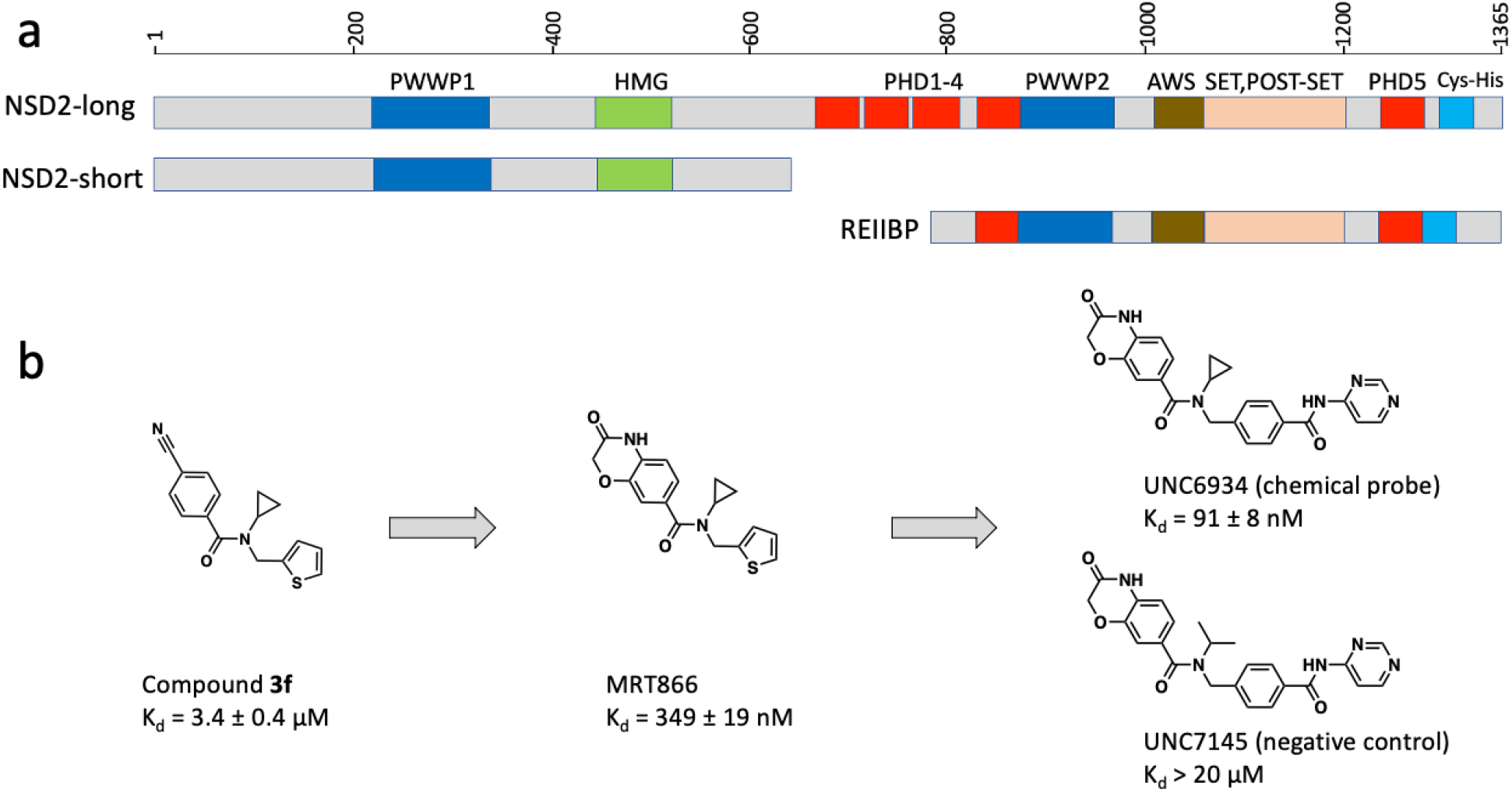
NSD2 protein architecture and NSD2-PWWP1 ligands. **(a)** Domain architecture of the three splicing isoforms, NSD2-long, NSD2-short, and REIIBP **(b)** Chemical series progression: successive rounds of optimization starting from compound **3f** ^19^ led to MRT866 and finally the chemical probe **UNC6934. UNC7145** is a structurally similar negative control compound.

Chromatin association is preserved with the PHD2 mutant but lost upon combined truncation of PWWP1, PHD1, PHD2 and part of PHD3 (i.e. the RE-IIBP isoform) due to cytoplasmic retention^14^. Conversely, nucleolar accumulation was observed in the case of NSD2-PWWP1 truncated variants^15,16^. Overall, these data suggest a complex regulatory system for NSD2 in which distinct combinations of structural modules contribute to subcellular localization, substrate engagement, and lysine methylation.

Due to their role in disease and biology, there has been much recent interest in targeting the NSD family of methyltransferases with small molecules. High-quality, cell-active, and selective inhibitors of NSD2 and NSD3 catalytic activity remain elusive; however, irreversible small molecule inhibitors of the NSD1 SET domain that demonstrate on target activity in NUP98-NSD1 leukemia cells have recently been reported^17^. We have had recent success targeting the PWWP domains of NSD3 and NSD2 suggesting that PWWP domains are druggable, while the PHD domains have so far not been targetable. Specifically, we reported a potent chemical probe targeting the N-terminal PWWP domain of NSD3 that repressed MYC mRNA levels and reduced the proliferation of leukemia cell lines^18^. Additionally, we recently described the development of the first antagonist of NSD2-PWWP1 which binds with modest potency and abrogates H3K36me2 binding^19^. Therefore, we hypothesized that targeting the PWWP domain(s) of NSD2 with highly potent and selective chemical probes may be a strategy to modulate NSD2 engagement with chromatin, subcellular localization, and/or catalytic function.

In this study, we report a first-in-class chemical probe, **UNC6934**, that selectively binds in the aromatic cage of NSD2-PWWP1, thereby disrupting its interaction with H3K36me2 nucleosomes. **UNC6934** potently and selectively binds full-length NSD2 in cells and induces partial disengagement from chromatin, consistent with a cooperative chromatin binding mechanism relying on multiple protein interfaces. **UNC6934** promotes nucleolar localization of NSD2, phenocopying previously characterized PWWP1-disrupting mutations prevalent in t(4;14) multiple myelomas^15,16^. Furthermore, we identified two active nucleolar localization sequences in NSD2 and demonstrated cooperativity between multiple chromatin reader modules to prevent nucleolar sequestration. Our data demonstrate that **UNC6934** is a potent and selective drug-like molecule suitable as a high-quality chemical probe to interrogate the function of NSD2-PWWP1.

## Results

### Discovery of a potent ligand targeting NSD2-PWWP1

We recently described the use of virtual screening, target class screening, and ligand-based scaffold hopping approaches to identify ligands of the NSD2 PWWP domains as starting points for further development^19^. This initial effort led to compound **3f** which binds NSD2-PWWP1 with a K_d_ of 3.4 ± 0.4 μM as determined by surface plasmon resonance (SPR). Based on the crystal structure of **3f** in complex with NSD2 (PDB 6UE6) molecular docking simulations predicted that a benzoxazinone bicyclic ring would favorably replace the cyanophenyl group of **3f**. We confirmed that compound **MRT866** binds NSD2-PWWP1 with a K_d_ of 349 ± 19 nM (**Fig. 1b, Supplementary Fig. 1**) and occupies the aromatic cage of PWWP1 similarly to compound **3f** as determined by x-ray crystallography (PDB 7LMT, **Supplementary Fig.1** and **Supplementary Table 1**). The benzoxazinone ring of **MRT866** makes more extensive Van der Waals interactions with NSD2-PWWP1 than **3f** and engages in an additional hydrogen bond with the side-chain of Q321. Further structure-based optimization focused on the replacement of the thiophene ring and resulted in **UNC6934** which binds NSD2-PWWP1 with a K_d_ of 91 ± 8 nM by SPR (**Fig. 1b and 2a**). Interestingly, conversion of the cyclopropyl group of **UNC6934** to an isopropyl moiety as in **UNC7145** (**Fig. 1b**) resulted in no appreciable binding up to 20 μM and therefore **UNC7145** is an ideal negative control compound. To confirm that **UNC6934** is binding in the methyl-lysine binding pocket of NSD2-PWWP1, we generated NSD2-PWWP1 with a key aromatic cage mutant (F266A). In contrast to wild type protein, **UNC6934** did not result in a significant thermal stabilization of the NSD2-PWWP1 F266A mutant (**Supplementary Fig. 2**). Furthermore, **UNC6934** is selective for NSD2-PWWP1 over 14 other human PWWP domains as assessed by differential scanning fluorimetry (DSF) (**Fig. 2b**) and did not inhibit any of a panel of 33 methyltransferase domains including the H3K36 methyltransferases, NSD1, NSD2, NSD3, and SETD2 (**Supplementary Fig. 3**). **UNC7145** was similarly inactive against all PWWP domains and methyltransferases tested (**Fig. 2a, Supplementary Fig. 3**).

**Figure 2:**
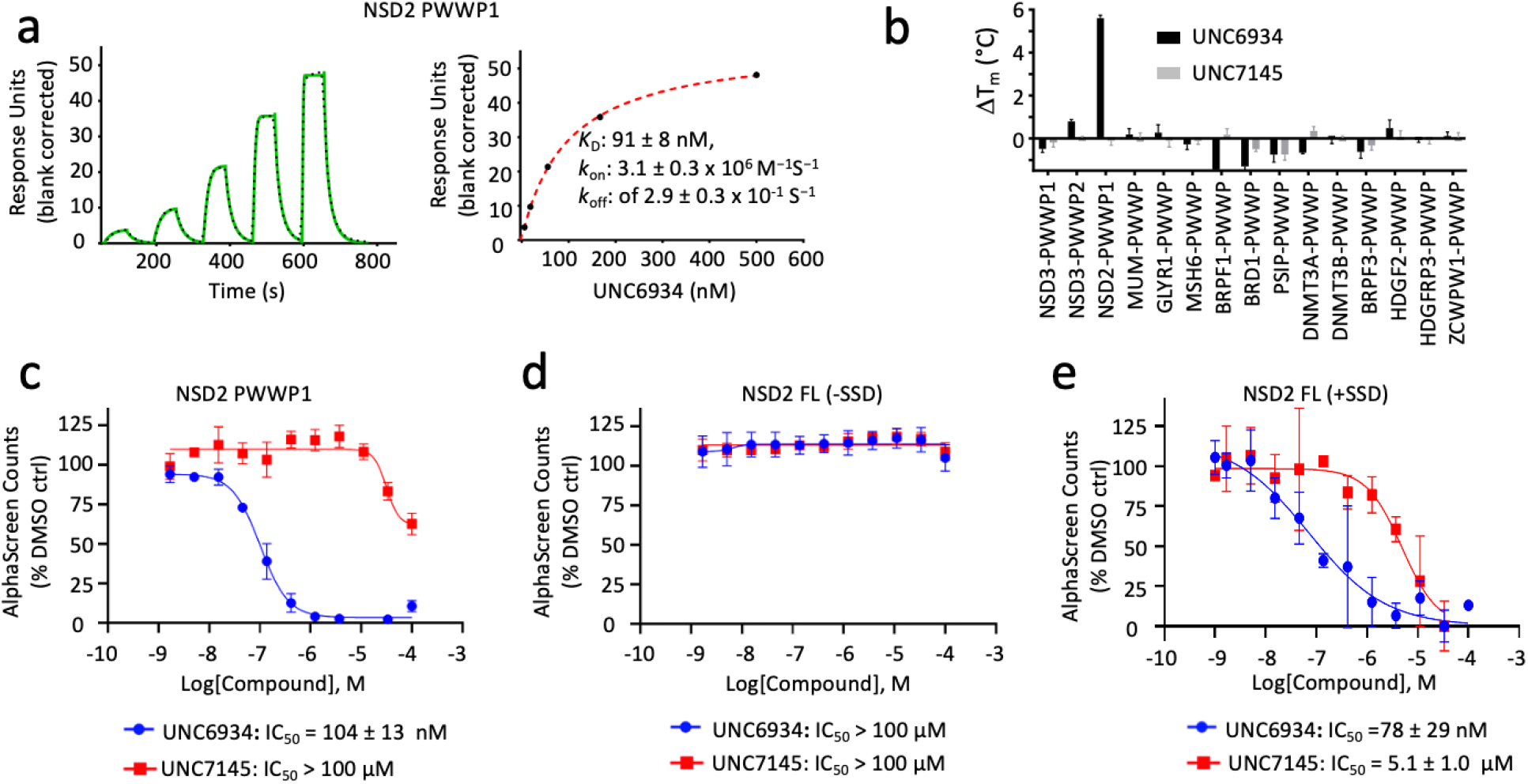
Biophysical characterization of UNC6934. **(a)** SPR analysis of the binding of **UNC6934** to NSD2-PWWP1. Left: Representative sensorgram (green) with the kinetic fit (black dots). Right: Steadystate response (black circles) with the steady-state 1:1 binding model fitting (red dashed line). Kinetic data indicated were obtained from triplicate experiments. **(b)** Selectivity profile of **UNC6934** and **UNC7145** against a panel of PWWP domains measured by DSF (compounds were tested at 100 μM). **(c)** Effect of **UNC6934** and **UNC7145** on the binding of NSD2-PWWP1 to nucleosomal H3K36me2 measured by AlphaScreen. Each binding interaction was performed in duplicate and all data normalized to the DMSO control. **(d, e)** Effect of **UNC6934** and **UNC7145** on the binding of full-length NSD2 to nucleosomal H3K36me2 measured by AlphaScreen in the absence (**d**) or presence (**e**) of salmon sperm DNA (SSD, 2 μg/ml).

### UNC6934 disengages NSD2-PWWP1 from methylated nucleosomes

NSD2-PWWP1 is postulated to stabilize the binding of NSD2 on chromatin, primarily through the recognition of H3K36me2^8^. Therefore, we next used an AlphaScreen-based proximity assay to investigate the effect of **UNC6934** on the interaction of NSD2 with recombinant semi-synthetic designer nucleosomes (dNucs)^9^. We first evaluated His-tagged NSD2-PWWP1 binding to a range of lysine-methylated semi-synthetic dNucs (me0, me1, me2, and me3 at H3K4, H3K9, H3K27, H3K36, and H4K20), and confirmed that NSD2-PWWP1 binds di- and tri-methyl H3K36 with a preference for the former (**Supplementary Fig. 4**), as previously reported^8^. **UNC6934** disrupted the interaction between NSD2-PWWP1 and nucleosomal H3K36me2 in a dose-dependent manner with an IC_50_ of 104 ± 13 nM, while **UNC7145** had no measurable effect (**Fig. 2c**).

To determine whether the PWWP1 domain is necessary for mediating the interaction between full-length NSD2 (fl-NSD2) and nucleosomes, we similarly tested the effect of **UNC6934** on fl-NSD2 binding to nucleosomal H3K36me2 in the AlphaScreen assay. Unlike with NSD2-PWWP1, we found that UNC6934 was unable to disrupt the interaction between fl-NSD2 and nucleosomal H3K36me2 (**Fig. 2d**). With H3K36me2 being proximal to nucleosomal DNA, we reasoned that electrostatic interactions between fl-NSD2 and DNA may abrogate disengagement of the protein by **UNC6934**. To test this hypothesis, we repeated the experiment in the presence of an excess of salmon sperm DNA (SSD), which is a commonly used blocker of non-specific DNA interactions. Under these conditions, **UNC6934** could effectively disengage fl-NSD2 from nucleosomal H3K36me2 (IC_50_ = 78 ± 29 nM) (**Fig. 2e**). Together, these results indicate that NSD2 binding to nucleosomes is multivalent, and therefore **UNC6934** can disengage fl-NSD2 from H3K36me2-modified nucleosomes only in the presence of excess competitive DNA.

### UNC6934 occupies the H3K36me2 binding pocket of NSD2-PWWP1

PWWP domains have both a methyl-lysine-binding pocket and a DNA-binding surface^12^. To better understand these interactions, we solved the crystal structures of NSD2-PWWP1 in complex with **UNC6934** (Supplementary Table 1). While DNA binds a basic surface area where the side-chains of K304, K309 and K312 are engaged in direct electrostatic interactions with the DNA phosphate backbone (**Fig. 3a**, PDB 5VC8), **UNC6934** occupies the canonical methyl-lysine-binding pocket adjacent to the DNA binding surface in an arrangement where the cyclopropyl ring is deeply inserted in the aromatic cage composed of Y233, W236 and F266 (**Fig. 3b, c**; PDB 6XCG). The extremely tight fit at the aromatic cage rationalizes the lack of binding of the corresponding negative control, **UNC7145**, where a bulkier isopropyl replaces the cyclopropyl group (**Fig. 1b**). The non-bonded carbons of an isopropyl group are separated by ~2.5 Å (ex: PDB 1NA3), while the corresponding atoms are 1.5Å apart in the cyclopropyl ring of **UNC6934**. The structure further explains the observed loss of binding of **UNC6934** to the F266A NSD2-PWWP1 mutant (**Supplementary Fig. 2**). Unlike the DNA binding surface, the **UNC6934** binding pocket is mildly electronegative, as would be expected for a site accommodating a positively charged methyl-lysine side-chain. Interestingly, the pocket is partly occluded in the apo structure and undergoes a conformational rearrangement of three side-chains (Y233, F266, E272) upon **UNC6934** binding (**Fig. 3d**). Overall, our structural data confirms that **UNC6934** competes directly with H3K36me2 for binding to NSD2-PWWP1 to disrupt high-affinity binding to H3K36me2 nucleosomes.

**Figure 3:**
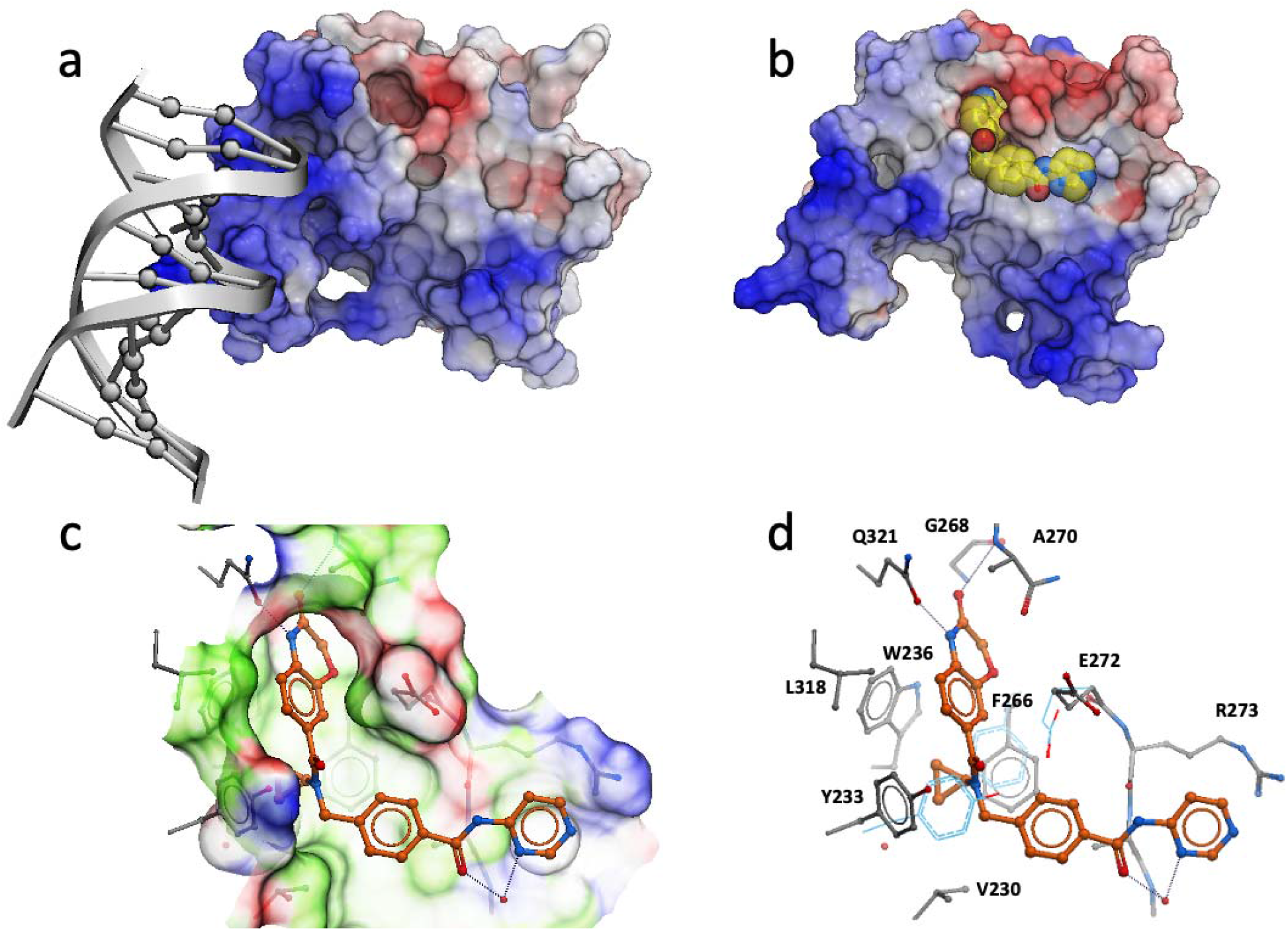
Crystal structures of NSD2 PWWP1. Structure of NSD2 PWWP1 in complex with (**a**) DNA [PDB code: 5VC8] and (**b**) **UNC6934** [PDB code: 6XCG]. Color coding according to electrostatic potential. Blue: electropositive. Red: electronegative. (**c,d**) Atomic interactions between **UNC6934** and surrounding side-chains. Color coding according to binding properties. Green: hydrophobic. Blue: hydrogen-bond donor. Red: hydrogen-bond acceptor. Side-chain conformations observed in the apo state are in light blue (**d**).

Our structural data also helps to explain the exquisite selectivity of **UNC6934** for NSD2-PWWP1 over other PWWP domains. Mapping the side-chains positioned within 5 Å of the bound ligand in our cocrystal structure onto a multiple sequence alignment of all human PWWP domains identified the degree of conservation among binding pocket residues exploited by **UNC6934**. We find that the binding pocket of NSD3-PWWP1 is by far the closest to that of NSD2-PWWP1, as only three of the side-chains lining the binding pocket are not conserved between the two proteins (**Supplementary Fig. 5**). In NSD2, G268 is at the bottom of a cavity accommodating the benzoxazinone of **UNC6934** and replacing this residue with the serine of NSD3 would be expected to occlude ligand binding. This is consistent with the ability of **UNC6934** to stabilize NSD2-PWWP1, but not NSD3-PWWP1 or any of the other PWWP domains in a thermal shift assay (**Fig. 2b**). These data strongly suggest that **UNC6934** is likely selective for NSD2-PWWP1 versus any other human PWWP domain. Further supporting the overall selectivity of the compound, **UNC6934** and **UNC7145** were profiled against a set of 90 central nervous system receptors, channels, and transporters (**Supplementary Table 2**). Of those that were inhibited by **UNC6934** greater than 50% at 10 μM (2 in total), the human sodium-dependent serotonin transporter receptor was the only protein inhibited by **UNC6934** with a measurable inhibitory constant (Ki = 1.4 ± 0.8 μM).

### UNC6934 selectively engages NSD2-PWWP1 in cells

To profile ligand selectivity and target engagement in a cellular context, we synthesized **UNC7096**, a biotin-labeled affinity reagent containing a close analog of **UNC6934** for chemical pulldown experiments (**Fig. 4a**). We first verified that **UNC7096** is a high-affinity NSD2-PWWP ligand by SPR, measuring a K_d_ of 46 nM, comparable to **UNC6934** (**Supplementary Fig. 6**). **UNC7096** efficiently enriched both NSD2-Short (MMSETI) and NSD2-Long (MMSETII) isoforms from KMS11 whole cell lysates as determined by western blotting (**Fig. 4b**). Chemiprecipitation of NSD2 by **UNC7096** could also be blocked by pre-incubation of KMS11 lysates with 20 μM of **UNC6934**, but not the negative control compound **UNC7145** (**Fig. 4b**). Label-free proteomic analysis of the **UNC7096** pulldown experiments identified NSD2 as the only protein significantly depleted by competition with **UNC6934** (**Fig. 4c**), whereas pre-incubation of lysates with **UNC7145** did not considerably alter the enrichment profile. Collectively, these experiments demonstrate selective binding of **UNC6934** to endogenous NSD2 protein within cell lysates.

**Figure 4:**
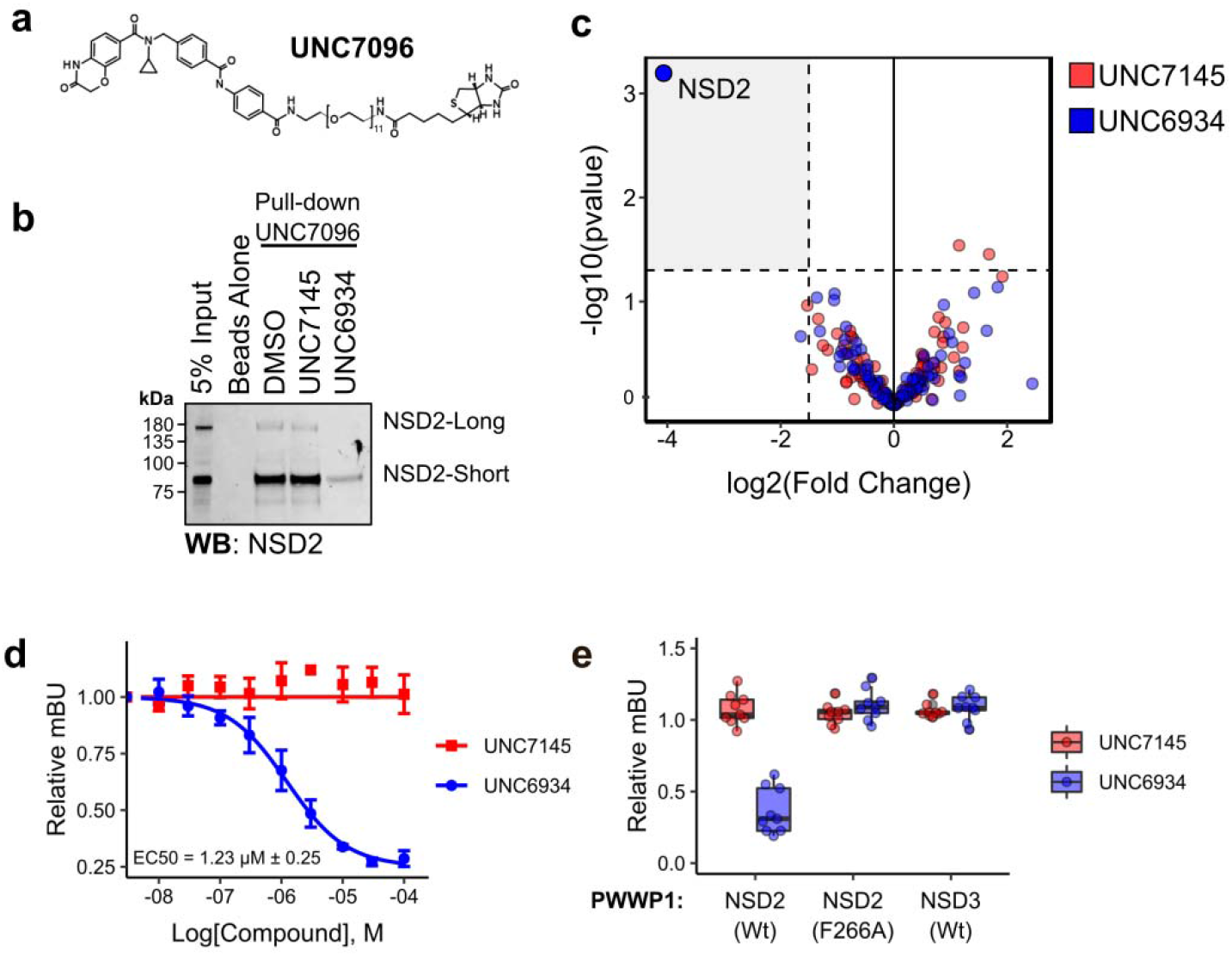
Cellular target engagement and selectivity. **(a)** Chemical structure of **UNC6934**-derived biotin affinity purification reagent, **UNC7096**. (**b**) NSD2 western blot analysis of **UNC7096** streptavidin pulldowns from KMS11 whole cell lysates after pre-incubation with either DMSO, 20 μM **UNC7175** or 20 μM **UNC6934**. (**c**) Selectivity profiling by chemical proteomics. Volcano plots show significantly displaced proteins from immobilized **UNC7096** pulldowns by competition with 20 μM **UNC6934 or UNC7145** relative to DMSO control. Significantly depleted hits were considered as fold enrichment less than −2 and a moderated p-value of 0.05 (n = 3). (**d**) Effect of **UNC6934** and **UNC7145** treatment on the nanoBRET signal produced by the interaction between NSD2-PWWP1-NanoLuc and Histone H3.3-Halo Tag in U2OS cells, EC_50_ = 1.23 μM ± 0.25 derived from three biological replicates (n=3). (**e**) Box and point plots showing the effect of 10 μM **UNC6934** or **UNC7145** on the interaction between H3.3-Halo Tag and NanoLuc-tagged wild-type NSD2-PWWP1, NSD2-PWWP1 F266A (aromatic cage mutant), or NSD3-PWWP1, using a nanoBRET protein-protein interaction assay (each point represents one technical replicate from three independent experiments (n =3).

To quantify the extent to which **UNC6934** engages NSD2 in live cells, we used a nanoBRET proteinprotein interaction assay to measure bioluminescence resonance energy transfer (BRET) between an Nluc-NSD2-PWWP1 fusion protein (donor) and a chloroalkane fluorophore (acceptor) bound to a Halotag-Histone H3.3 fusion in U2OS cells^20^. We observed a dose-dependent decrease in the BRET signal upon treatment with **UNC6934** (EC_50_ = 1.23 ± 0.25 μM), but not with the negative control compound **UNC7145** (**Fig. 4d**). The BRET signals of an NSD2-PWWP1 aromatic cage mutant (F266A) or NSD3-PWWP1 were unaffected by **UNC6934** (10 μM), supporting selective engagement of the NSD2-PWWP1 methyl-lysine binding pocket (**Fig. 4e**). Further supporting minimal off-target activity and the suitability of **UNC6934** to query NSD2-PWWP1 function in cells, we found no acute cytotoxic effects upon treatment of five common cell lines with **UNC6934** or **UNC7145** up to 10 μM over 72 hours (**Supplementary Fig. 7**).

### Perturbation of NSD2’s Chromatin Binding Domains Promotes Localization to the Nucleolus

Reader domains are critical for the recruitment and positioning of epigenetic proteins at defined loci across the genome and small molecule antagonism of reader domains is known to alter the localization of chromatin-associated target proteins^21,22^. We therefore reasoned that PWWP1 antagonism by **UNC6934** may affect NSD2 localization within the nucleus. To test this hypothesis, we used confocal microscopy to evaluate the localization of endogenous NSD2 in U2OS cells treated for four hours with 5 μM **UNC6934** or negative control **UNC7145** (**Supplementary Fig. 8)**. Upon treatment with **UNC6934**, we observed an increase in NSD2 signal within nucleoli, the sub-nuclear membrane-less organelles that house ribosomal DNA for pre-ribosomal transcription and processing. Interestingly, t(4;14) chromosome translocations in multiple myeloma, which juxtapose the IgH enhancer and NSD2 promoting overexpression of NSD2, can result in truncation/inactivation of PWWP1 and nucleolar enrichment, suggesting that the PWWP1 domain contributes to the exclusion of NSD2 from the nucleolus^15,16^. Therefore, **UNC6934** appears to phenocopy NSD2 N-terminal PWWP1 truncations while **UNC7145** has no effect.

To validate this result, we repeated the imaging experiments while co-staining for the nucleolar marker fibrillarin and measured the extent of NSD2 and fibrillarin co-localization using Pearson correlation (**Fig. 5a,b**). The Pearson correlation coefficient (PCC) is a common statistic used to describe co-localization; it measures the correlation in signal intensity between two fluorescent molecules (values range between −1 and 1 representing no correlation and absolute correlation, respectively). We observed a significant increase in the correlation between NSD2 and fibrillarin signal in response to **UNC6934**, confirming an increase in the nucleolar localization of NSD2. These data support the engagement of endogenous NSD2 by **UNC6934** in cells and suggest that loss of H3K36me2 binding by NSD2-PWWP1 promotes nucleolar accumulation of NSD2. While we found no significant effect on ribosomal RNA transcription in response to **UNC6934** (**Supplementary Fig. 9**), we do observe a steady-state pool of nucleolar NSD2 that is sensitive to RNA polymerase I inhibition (actinomycin D; 50 nM) or genotoxic agents (doxorubicin; 1 μM), conditions known to significantly alter the protein composition of nucleoli^23,24^ (**Supplementary Fig. 10**). Overall, these results indicate that **UNC6934** mediated antagonism of PWWP1 leads to accumulation of endogenous NSD2 in the nucleolus.

**Figure 5:**
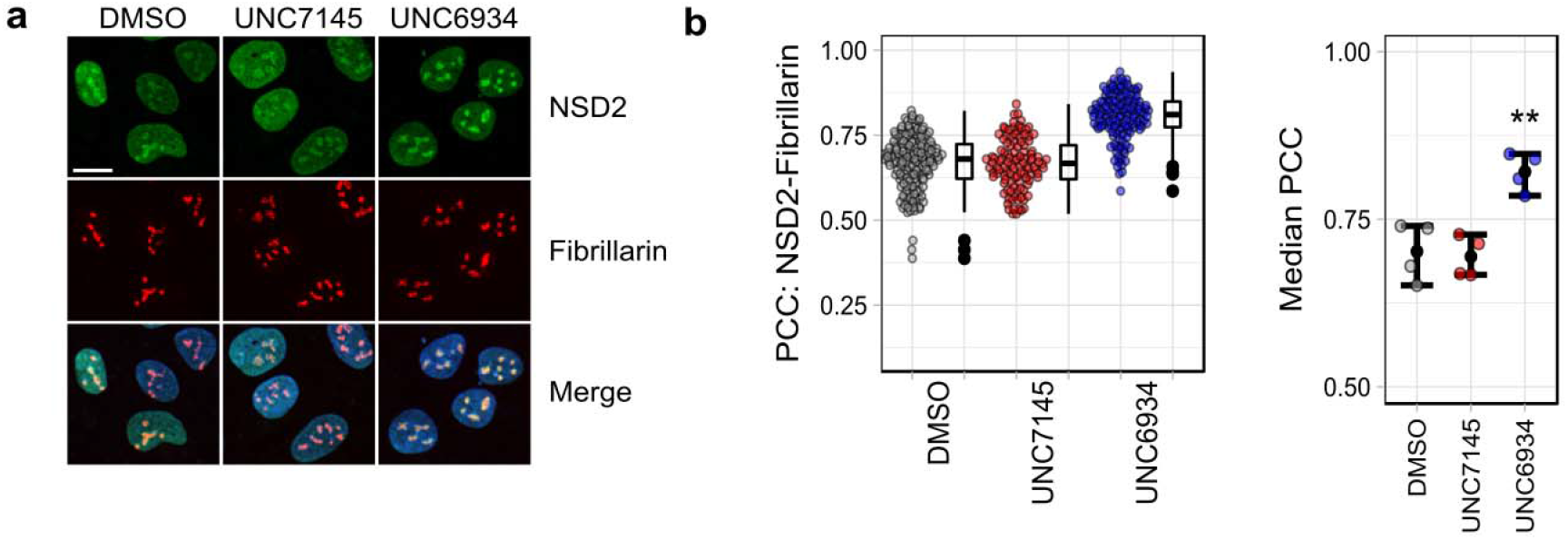
UNC6934 promotes enrichment of endogenous NSD2 in the nucleolus. (**a**) Representative confocal microscopy images of NSD2 and fibrillarin staining in U2OS cells treated for 4 hours with DMSO vehicle control, 5 μM **UNC7145**, or 5 μM **UNC6934** (scale bar = 15 μm). (**b**) Quantification of co-localization between NSD2 and the nucleolar marker fibrillarin as determined by Pearson correlation coefficient (PCC). On the left a representative box and point plot shows the data for one biological replicate with each point representing the PCC per nuclei measured across at least four fields of view. On the right, the median PCC is shown for four independent biological replicates (n = 4, ** p-value = 0.0055 Welch unpaired t-test compared to DMSO control).

To test if subnuclear localization is exclusively mediated by the PWWP1 domain, we mutated GFP-tagged NSD2 at several critical sites within distinct chromatin-recruitment modules. This included an N-terminal short linear motif that engages BET proteins (K125A)^21,25^, two key aromatic cage residues in the PWWP1 domain (W236A and F266A)^12^, two PHD2 mutants that disrupt recruitment to target loci and disable H3K36me2 methyltransferase activity in cells (H762R and H762Y)^14^, a presumptive inactivating aromatic cage mutation in PWWP2 (W894A), and a catalytic-dead mutant of the methyltransferase domain (Y1092A)^1^. We found that disruption of any one of the canonical reader domains (PWWP1, PHD2, and PWWP2) promoted enrichment of NSD2 in nucleoli (**Fig. 6a, b**). These observations suggest that it is the loss of chromatin binding that leads to the nucleolar retention of NSD2, and that multiple NSD2 reader domains cooperate to maintain appropriate nuclear sublocalization.

**Figure 6:**
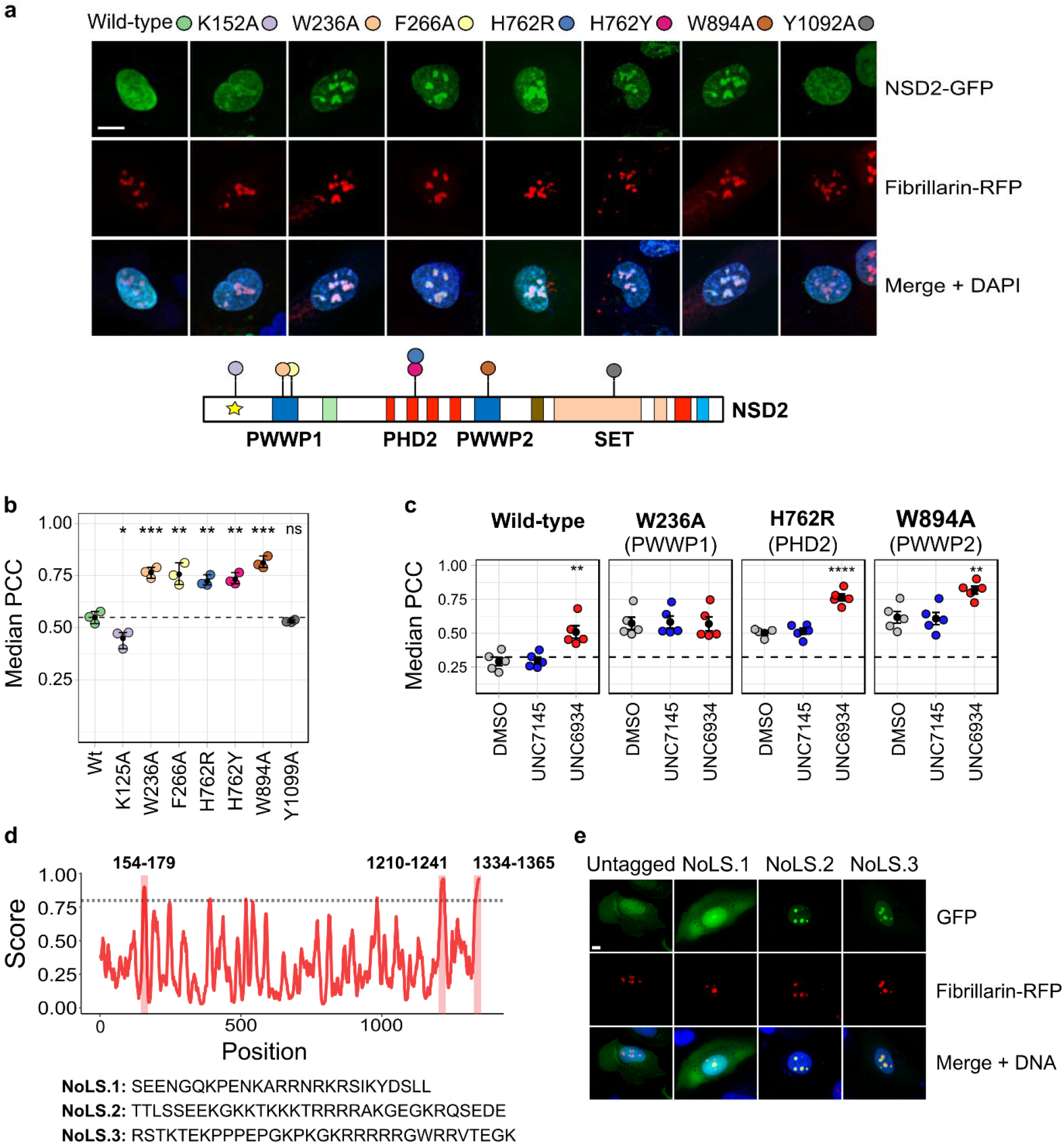
Multivalent interactions dictate the nuclear localization of NSD2. (**a**) Representative confocal microscopy of U2OS cells co-expressing GFP-tagged NSD2 wild-type/mutants and RFP-tagged fibrillarin (scale bar = 15 μm). Domain schematic of NSD2 indicates the position of each mutation. (**b**) Quantification of co-localization between RFP-tagged fibrillarin and NSD2 wild-type or mutants by median Pearson correlation coefficient (PCC) per biological replicate between NSD2 and fibrillarin signal across three independent experiments (n=3; p-values derived from a Welch’s unpaired t-test compared to Wt; K125A = 0.036, W236A = 0.0001, F266A = 0.0084, H762R = 0.0016, H762Y = 0.0014, W894A = 0.0004, Y1099A = 0.38). For each experiment, PCC/nuclei was measured across sixteen fields of view. (**c**) Assessing domain cooperativity by treating NSD2-GFP point mutants with DMSO control, 5 μM **UNC7145** or 5 μM **UNC6934** and measuring co-localization by PCC (n=5, significant p-values derived from a Welch’s unpaired t-test compared to the DMSO control for each panel are indicated in order as ** = 0.0059, **** = 8.7 x 10^-5^, ** = 0.0062). **(d)** Computational prediction of Nucleolar Localization Sequences in NSD2 using the Nucleolar localization sequence Detector algorithm (NOD)^26^. (**e**) Representative fluorescent images of cells expressing GFP tagged with putative nucleolar localization sequences from NOD.

We next used **UNC6934** to test the cooperativity of the NSD2 reader domains towards nucleolar localization. To do so, we treated cells transfected with RFP-fibrillarin and GFP-tagged NSD2 (wild-type, W236A, H762R, or W894A) for four hours with 5 μM **UNC6934** or **UNC7145**. In cells expressing wildtype NSD2-GFP we observed an increase in nucleolar signal upon treatment with **UNC6934**. Importantly, while the PWWP1 aromatic cage mutant (W326A) had a higher baseline nucleolar localization compared to WT, no further increase was observed upon **UNC6934** treatment, again supporting PWWP1-dependent activity of the probe (**Fig. 6c**). However, in cells expressing the PHD2 and PWWP2 mutants, we observed an additive effect with **UNC6934** treatment. In addition to higher baseline nucleolar localization due to their respective mutations, there was a further increase in nucleolar colocalization upon **UNC6934** treatment compared to both the DMSO control and the negative control **UNC7145**. These observations support a model in which NSD2 reader domains act cooperatively in recruiting NSD2 to chromatin and preventing nucleolar sequestration. Antagonizing the interaction between NSD2-PWWP1 and H3K36me2 may therefore not be sufficient to fully disengage full-length NSD2 from chromatin. Indeed, we only observed a modest release of full-length NSD2 from chromatin upon treatment with **UNC6934** in cell fractionation experiments, as well as no changes in global H3K36me2 levels or the proliferation of KMS11 t(4;14) multiple myeloma cells grown on bone marrow stroma in response to **UNC6934** (**Supplementary Fig. 11**).

To further define the features of NSD2 that drive its nucleolar targeting, we used NoD (Nucleolar localization sequence Detector)^27^ to computationally predict putative nucleolar localization sequences (NoLS) within NSD2 (**Fig. 6d**). Of the three NoLS predicted with high-confidence, we found that two were arginine-rich sequences within the C-terminus that could robustly target GFP to nucleoli (**Fig. 6e**). These results suggest a competitive situation between chromatin reader-domains and NoLS sequences, and support a model in which perturbation of NSD2 chromatin-binding modules enables the activity of C-terminal NoLS’s to dominate, leading to nucleolar accumulation.

## Discussion

Here we describe the discovery of **UNC6934**, the first chemical probe to target NSD2, a potent and well-characterized driver of both hematological malignancies and solid tumours. **UNC6934** follows the recent discovery of **BI-9321**, which targets the closely-related PWWP1 domain of NSD3^18^. **UNC6934** and **BI-9321** are chemically distinct and selective, establishing PWWP domains as tractable targets for future chemical biology and drug discovery efforts.

Importantly, **UNC6934** satisfies stringent criteria for high-quality chemical probes as fit-for-purpose research tools^28-30^, including <100 nM *in vitro* potency (K_d_ of 91 ± 8 nM by SPR; IC_50_ of 104 ± 13 nM by Alphascreen) and significant cellular activity at 1 μM (IC_50_ = 1.23 ± 0.25 μM by nanoBRET PPI assay). Additionally, we found no significant off-targets by chemical proteomics and *in vitro* screening of functionally relevant proteins, including a large number of human PWWP domains, methyltransferases, and membrane proteins. Further demonstration of cellular activity and specific target engagement is evident from the robust changes in endogenous NSD2 localization in response to PWWP1 antagonism by **UNC6934**. We also show limited cytotoxicity by **UNC6934** and its negative control counterpart **UNC7145**, signifying their suitability for cell biology experiments exploring the function of the NSD2 PWWP1 reader domain.

Nucleoli are dynamic membrane-less nuclear structures that not only act as the site of ribosome transcription and pre-assembly, but also as integral organizational hubs in the regulation of many non-canonical functions^31^. It is now widely appreciated that the shuttling of proteins between the nucleolus and nucleoplasm is a critical feature of nuclear biology, regulating many processes, including stress response, DNA repair, recombination, and transcription^31-33^. Here we define active nucleolar localization sequences in the NSD2 C-terminus, which likely drive nucleolar sequestration of NSD2 in response to modulation of its chromatin-binding domains. Altering the balance of nucleoplasmic versus nucleolar NSD2 through PWWP1 antagonism by **UNC6934** did not have a significant effect on ribosome transcription or global levels of H3K36me2, suggesting instead that these features may provide a mechanism to rapidly tune the sub-nuclear localization of NSD2 in response to stimuli. Supporting this idea, a recent report showed that epigenetic proteins, including NSD2, are sequestered within the nucleolus in response to heat shock stress as a mechanism for subsequent rapid recovery and epigenome maintenance (data from Azkanaz et al. highlighting NSD2 shown as **Supplementary Fig. 12**)^34^. Given our observations of a steady-state pool of nucleolar NSD2, the question remains how balance and control of nucleolar-nucleoplasmic NSD2 levels may influence NSD2 function in normal and disease biology. Cytoplasmic localization of an NSD2 variant lacking PWWP1 and the first 3 PHD domains was also reported^14^, suggesting that the sub-cellular compartmentalization of NSD2 is fine-tuned by its reader domains, which could be achieved by masking sub-cellular localization sequences or by engagement of subcellular-specific substrates.

Our data highlight the multivalent nature of NSD2 recruitment to chromatin, whereby NSD2 chromatin reader domains and DNA binding interfaces act cooperatively to coordinate its activity on chromatin. These findings highlight the utility of **UNC6934** as a tool to interrogate the contributions of NSD2-PWWP1 in the interplay between reader domains. Finally, because of its role in multiple myeloma and other cancers, NSD2 has long been a drug target of interest, but despite much community effort, there is no selective, cell-active inhibitor of its catalytic activity. **UNC6934** provides a clear starting point for the development of bifunctional molecules, like PROTACs, able to induce proteasomal degradation of NSD2 to antagonize its function in disease.

## Acknowledgements

The Structural Genomics Consortium is a registered charity (no: 1097737) that receives funds from; AbbVie, Bayer AG, Boehringer Ingelheim, Canada Foundation for Innovation, Eshelman Institute for Innovation, Genentech, Genome Canada through Ontario Genomics Institute [OGI-196], EU/EFPIA/OICR/McGill/KTH/Diamond Innovative Medicines Initiative 2 Joint Undertaking [EUbOPEN grant 875510], Janssen, Merck KGaA (aka EMD in Canada and US), Merck & Co (aka MSD outside Canada and US), Pfizer, Takeda and Wellcome [106169/ZZ14/Z]. We acknowledge the Natural Sciences and Engineering Research Council of Canada (NSERC) for a postdoctoral fellowship awarded to DD. This work was supported by the National Cancer Institute, NIH (grant R01CA242305) to L.I.J. MS gratefully acknowledge NSERC (grant RGPIN-2019-04416). Research in EpiCypher was supported by NIH grants R44GM117683 and R44GM116584. We thank Taraneh Hajian for purfying proteins. We thank the University of North Carolina’s Department of Chemistry Mass Spectrometry Core Laboratory, especially Diane Wallace, for their assistance with mass spectrometry analysis. The Mass Spectrometry Core Laboratory is supported by the National Science Foundation under Grant No. (CHE-1726291). Research reported in this publication was supported in part with funding by the University of North Carolina’s School of Medicine Office of Research. We thank Frances Potjewyd for reviewing the primary synthesis data supporting this manuscript. Receptors, channels and transporters binding profiles were generously provided by the National Institute of Mental Health’s Psychoactive Drug Screening Program, Contract # HHSN-271-2018-00023-C (NIMH PDSP). The NIMH PDSP is Directed by Bryan L. Roth at the University of North Carolina at Chapel Hill and Project Officer Jamie Driscoll at NIMH, Bethesda MD, USA. For experimental details please refer to the PDSP web site https://pdsp.unc.edu/ims/investigator/web/.

## Author Contributions

Experimental design: DD, RPH, RFF, AAH, MZ, MRM, SA, EM, MMS, DK, MCK, JM, PJB, MV, CHA, DBL, LIJ, MS; data generation: DD, RPH, RFF, AAH, MZ, NM, MRM, SA EM, FL, IC, AB, SQ, ML YL, MMS, AD, SK, TA, IKP, NWH, MJM, MAC, EG; data analysis: DD, RPH, RFF, AAH, MZ, MRM, SA, EM, MMS, IKP, NWH, MAC, DK, MCK, JM, PJB, MV, CHA, DBL, LIJ, MS; supervision: DK, JFG, MCK, JM, MV, CHA, DBL, LIJ, MS; manuscript writing: DD, RPH, MCK, MV, CHA, DBL, LIJ, MS; funding: CHA, LIJ; manuscript review: all.

## Methods

### Expression and Purification of biotinylated NSD2-PWWP1

#### Construct and Expression

DNA fragment encoding human NSD2 (residues 208-368) was amplified by PCR and sub-cloned into p28BIOH-LIC vector, downstream of an AviTag and the upstream of a poly-histidine coding region. Following transformation into *E. Coli* BL21 (DE3) the cells were amplified at 37°C by inoculating Terrific Broth with overnight culture, both supplemented with 50 μg/ml Kanamycin and 35 μg/ml chloramphenicol. When the OD600 of the culture reached 0.8-1.5, the temperature was lowered to 16°C and the target protein was over-expressed by inducing cells with 0.5 mM IPTG (isopropyl-1-thio-D-galactopyranoside) and D-Biotin was added at 10 μg/mL final concentration and incubated overnight before being harvested (7000 rpm for 10 min at 4 °C) using a Beckman Coulter centrifuge.

#### Harvest and cell lysis

Harvested cells were re-suspended in 20 mM Tris-HCl, pH 7.5, 500 mM NaCl, 5 mM imidazole and 5% glycerol, 1X protease inhibitor cocktail (100 X protease inhibitor stock in 70% ethanol (0.25 mg/ml Aprotinin, 0.25 mg/ml Leupeptin, 0.25 mg/ml Pepstatin A and 0.25 mg/ml E-64) or Pierce™ Protease Inhibitor Mini Tablets, EDTA-free. The cells were lysed chemically by rotating 30 min with 0.5% CHAPS, 1 mM DTT, 1 mM PMSF and 15 μL Benzonase Nuclease (In-House) followed by sonication at frequency of 7 (5” on/7” off) for 5 min (Sonicator 3000, Misoni). The crude extract was clarified by high-speed centrifugation (60 min at 36,000 ×g at 4 °C) by Beckman Coulter centrifuge.

#### Purification

The clarified lysate was then loaded onto an open column containing pre-equilibrated Ni-NTA (Qiagen). The column was washed with 20 mM Tris-HCl, pH 7.5, 500 mM NaCl, 5% glycerol, 5 mM imidazole, then with 1 mM D-Biotin in PBS, followed by 20 mM Tris-HCl, pH 7.5, 500 mM NaCl, 5% glycerol, 20 mM imidazole. Finally, the protein was eluted by running 20 mM Tris-HCl, pH 7.5, 500 mM NaCl, 5% glycerol, 250 mM imidazole. The eluted protein was then supplemented with 1mM TCEP and concentrated to be further purified by gel filtration on a HiLoad Superdex200 16/600 using an ÄKTA Pure (GE Healthcare). The gel filtration column was pre-equilibrated with 20 mM Tris-HCl, pH 8.0, 500 mM NaCl, 5% glycerol. The purity of the fractions was assessed on SDS-PAGE gels and pure fractions (>95%) were pooled, concentrated and flash frozen.

### Surface Plasmon Resonance (SPR)

SPR was performed as described previously [1] with modifications. Biotinylated NSD2-PWWP1 domain was immobilized on the flow cell of a SA sensor chip in 1x HBS-EP buffer (10 mM HEPES pH 7.4, 150 mM NaCl, 3 mM EDTA, 0.05% Tween-20), yielding approximately 5000 response units (RU) according to manufacturer’s protocol while another flow cell was left empty for reference subtraction. UNC6934 and UNC7145 were tested at 2 μM as the highest concentration and dilution factor of 0.33 was used to yield 5 concentrations. Experiments were performed using the same buffer with 0.5% DMSO in single cycle kinetic with 60s contact time and a dissociation time of 120s at a flow rate of 75 μL/min.

### Molecular Docking

The X-ray structure of the PWWP domain of NSD2 in complex with **3f** (PDB ID: 6UE6) was prepared with PrepWizard (Schrodinger, New York) using the standard protocol, including the addition of hydrogens, the assignment of bond order, assessment of the correct protonation states, and a restrained minimization using the OPLS3 force field. Receptor grids were calculated at the centroid of the ligand with the option to dock ligands of similar size and a hydrogen bonding constraint with the backbone of A270 was defined.

Over 6,000 commercially available chemical analogs of **3f** were prepared with LigPrep (Schrodinger, New York). The resulting library was then docked using Glide SP (Schrodinger, New York) with default settings. Also, the core docking option was turned on to allow only ligand poses that have their core aligned within 1.0 Å of the reference core (the cyclopropyl and the amide group of **3f**). Only 448 compounds fitted and were ranked by Glide. Finally, after a visual inspection, 20 compounds were ordered. Optimization and SAR leading from MRT866 to UNC6934 was guided by free energy perturbation and will be presented elsewhere.

### dCypher binding assays

Recombinant semi-synthetic designer nucleosomes (dNucs) were from *EpiCypher*. Two phases of dCypher^®^ testing on the PerkinElmer AlphaScreen^®^ platform were performed as previously described^9^. In Phase A, HIS-tagged human NSD2 PWWP1 (aa208-368) was titrated to positive (H3K36me2) and negative (H3K36me0) control nucleosomes at a range of NaCl concentrations (50, 100, 150, 200 and 250mM) to determine binding curves and identify salt preference, level of sensitivity, signal saturation, and signal over background. In Phase B, an optimal HIS-NSD2 PWWP1 concentration was used to probe the biotinylated nucleosome panel (me0-1-2-3 at H3K4, H3K9, H3K27, H3K36 and H4K20). For all dCypher testing 5 μl of HIS-tagged NSD2 (format / concentration as indicated) was incubated with 5 μl of biotinylated nucleosome (10 nM) for 30 min at 23°C in salt-optimized assay buffer (150mM NaCl, 20mM Tris pH 7.5, 0.01% BSA, 0.01% NP-40, 1mM DTT]) in a 384-well Optiplate (*PerkinElmer*, 6007299). A mixture of 10 μl of 5 μg/ml Nickel-chelate acceptor beads (*PerkinElmer*) and 20 μg/ml streptavidin donor beads (*PerkinElmer*) was prepared in nucleosome bead buffer (as assay buffer minus DTT) and added to each well. The plate was incubated at room temperature in subdued lighting for 60 min and the AlphaScreen signal measured on a PerkinElmer 2104 EnVision (680-nm laser excitation, 570-nm emission filter ± 50-nm bandwidth). Each binding interaction was performed in duplicate.

When present, salmon sperm DNA (concentration as noted) was added at the NSD2-Nucleosome pre-incubation step. For inhibitor testing compound stocks were dissolved in DMSO and diluted to 400 μM in assay buffer (and further dilutions prepared in assay buffer supplemented with 2% DMSO). 5 μL of compound and 5 μL of NSD2 PWWP1 (40 nM) or NSD2 FL (10 nM) were combined and incubated for 15 minutes at 23°C. Then 5 μL of H3K36me2 dNuc (10 nM) in assay buffer was added and incubated for 30 minutes at 23°C. Donor/Acceptor beads and signal detection were as above. IC_50_ apparent was determined by a 10-point data curve (in duplicate) to identify upper and lower plateaus, with values calculated for compounds inhibiting ≥50% of signal.

### Selectivity assays

Selectivity of UNC6934 for NSD2-PWWP1 over 14 other PWWP domains was tested using differential scanning fluorimetry (DSF) as previously described^35^. In brief, the assay determined the effect of 100 μM of UNC6934 and UNC7145 on the thermal stability of NSD2-PWWP1 and 14 PWWP domains from various proteins. All proteins tested in these selectivity experiments were used at final concentration of 0.1 mg/ml in buffer (100mM HEPES, pH 7.5 and 150 mM NaCl) containing 5x Sypro Orange (Invitrogen, 5,000× stock solution). The temperature scan curves were fitted to a Boltzmann sigmoid function, and the T_m_ values were obtained from the midpoint of the transition. More than 2 °C increase in stability of the protein (ΔT_m_ > 2 °C) was considered as a confirmation of binding. Selectivity against 33 methyltransferases were determined as previously described^36^. Selectivity against 90 central nervous system receptors, channels and transporters was conducted at the NIMH Psychoactive Drug Screening Program^37^.

### Crystallography

#### Protein Expression and Purification

NSD2-PWWP1 (aa211-350) was subcloned into the pET28-MHL vector. The recombinant protein was overexpressed in *Escherichia coli* BL21 (DE3)-V2R-pRARE2 induced with 0.25mM isopropyl-D-thiogalactopyranoside (IPTG) at 16 °C overnight. The cell pellet was dissolved and further lysed in a buffer containing 20 mM Tris-HCl, pH 7.5, 500 mM NaCl, and 5 % glycerol. Supernatant was collected after centrifugation at 16,000 ×g for 1 hour. The supernatant was loaded on to a Ni-NTA (Ni2+-nitrilotriacetate) column(GE Healthcare) pre-equilibrated with lysis buffer, washed with 20 mM Tris-HCl, pH 7.5, 500 mM NaCl and 25 mM imidazole, and then eluted in 20 mM Tris-HCl, pH 7.5, 150 mM NaCl and 250 mM imidazole. Purified protein was treated by tobacco etch virus (TEV) proteases to remove the tag. The treated sample was further analyzed by affinity chromatography and gel-filtration column (GE Healthcare). Finally, the pure protein was concentrated to 20 mg/ml in a buffer containing 20 mM Tris-HCl, pH 7.5, and 150 mM NaCl.

#### Crystallization

The purified protein was mixed at a molar ratio of 1:3 with compound **UNC6934** or **MRT866** followed by incubation at room temperature for 30 minutes. The **UNC6934** protein mixture precipitated quickly when adding the compound into the protein solution. After centrifugation at 13,000×g for 10 min, the supernatant mixture was crystallized by the sitting drop vapor diffusion method at 18 °C by mixing 0.5μl of the complex samples with 0.5μl of the reservoir solution, and crystals appeared in a condition containing 1.6M (NH_4_)2SO_4_, 0.01M MgCl2 and 0.1M HEPES pH 7.5 after two days. The **MRT866** protein mixture was clear and crystallized using the sitting drop vapor diffusion method at 18 °C and a crystallization buffer of 25% P3350, 0.2M MgCl2, and 0.1M HERES (pH7.5). Prior to data collection, both kinds of crystals were freshly soaked in the mother liquor complemented with 15-20% glycerol and flash-frozen in liquid nitrogen.

#### Data Collection and Structure Determination

The diffraction data for NSD2-PWWP1+UNC6934 was collected at 100K on the home source Rigaku FR-E superbright and data set was processed using the HKL-3000 suite^38^. The structure was solved by molecular replacement methods with PHASER using PDB entry 5VC8 as search template. REFMAC was used for structure refinement^39^. GRADE (http://www.globalphasing.com) was used to generate all restrains for compound refinement. Graphics program COOT^40^ was used for model building and visualization. Molprobity^41^ was used for structure validation. X-ray diffraction data for NSD2-PWWP1+ MRT866 was collected at 100K on APS beamline 19ID (Argonne National Laboratory). The data set was processed using the XDS suite^42^. The structures was solved by molecular replacement with MOLREP^43^. Geometry restraints for compound MRT866 refinement were prepared with ACEDRG^44^, Graphics program COOT^40^ was used for model building and visualization. Restrained refinement and validation were conducted with BUSTER^45^, and MOLPROBITY^41^, respectively

### Plasmids

For NanoBRET PPI assays, the Histone H3.3-HaloTag^®^ Fusion Vector was purchased from Promega, while the sequences for NSD2 or NSD3 PWWP1 domain was cloned in-frame into a pNLF1-C [CMV/Hygro] vector from Promega using the In-Fusion HD Cloning kit (Takara). Point mutations were introduced using the Q5^®^ Site-Directed Mutagenesis Kit (New England Biolabs). For fluorescence microscopy, C-terminal GFPSpark^®^-tagged NSD2 wild-type was purchased from Sino Biological. NSD2 point mutants were generated using mutagenic primers with Q5^®^ Site-Directed Mutagenesis Kit from New England Biolabs. pEGFP-C1-Fibrillarin was a gift from Sui Huang (Addgene plasmid # 26673; RRID:Addgene_26673)^46^.

### Cell Culture

Cell lines were cultured according to standard aseptic mammalian tissue culture protocols in 5% CO_2_ at 37 °C with regular testing for mycoplasma contamination using the MycoAlert™ Mycoplasma Detection Kit (Lonza). HCT 116, HEK 293, HT1080, MCF7, and U2OS cells were cultured in DMEM supplemented with 10% fetal bovine serum (FBS) (Wisent) and 100U/100 μg/mL pencillin/streptomycin (Wisent). OP9 bone marrow stromal cells, prior to proliferation assays, were culture in α-MEM media with GlutaMAX (Invitrogen) containing 20% fetal bovine serum (FBS (Wisent), 55 μmol/L β-mercaptoethanol (Invitrogen) and 100U/100 μg/mL penicillin/streptomycin (Wisent). KMS-11 parental and TKO2 cells, a kind gift from Dr. Johnathan Licht, were cultured in RPMI (Wisent) supplemented with 10% FBS (Wisent) and 100U/100 μg/mL penicillin/streptomycin (Wisent).

### NanoBRET Protein-protein Interaction Assay

For NanoBRET protein-protein interaction assays, U2OS cells plated in six well plates were transfected with 1.8□μg of C-terminally tagged histone H3-HaloTag Fusion Vector DNA and 0.2□μg of C-terminally tagged NSD2-PWWP1-NanoLuc (Wt or F266A) or NSD3-PWW1-NanoLuc using X-tremeGENE HP DNA Transfection Reagent (Sigma), following the manufacturer’s instructions. The next day, cells were trypsinized and resuspended in DMEM/F12 (no phenol red) supplemented with 4% FBS, penicillin (100□U□ml ^1^) and streptomycin (100□μg□ml ^1^) at a density of 1.1□×□10^5^□cells/ml. Cells were then divided into two pools. HaloTag NanoBRET 618 Ligand (Promega) was added to the first pool and DMSO was added to the second pool, following the manufacturer’s instructions. Cells were plated in 96-well white plastic plates (Greiner) in the presence or absence of compounds for 20 □h. Next, NanoBRET NanoGlo Substrate (Promega) solution was added to each well. Donor emission at 450□nm (filter: 450□nm/BP 80□nm) and acceptor emission at 618□nm (filter: 610□nm/LP)) was measured within 10□min of substrate addition using a CLARIOstar microplate reader (Mandel). Mean corrected NanoBRET ratios (mBU) were calculated by subtracting the mean of 618/460 signal from cells without a NanoBRET 618 Ligand□×□1,000 from the mean of 618/460 signal from cells with a NanoBRET 618 Ligand□×□1,000.

### Chemical Proteomics

To prepare whole cell lysates, KMS11 cells were washed 2 times with 1x PBS, lysed by resuspension in high-salt lysis buffer (20 mM HEPES pH 7.5, 350 mM KCl, 1% Triton X-100 + a protease inhibitor cocktail containing aprotinin, leupeptin, pepstatin A, and E-64) and passed through a 25 gauge needle 5 times followed by a 20 min incubation on ice. Cell lysates were cleared by centrifugation at 18 000 x g for 20 minutes at 4°C. Cleared supernatant was diluted to 150 mM KCl and 0.4% Triton X-100 with 20mM HEPES pH7.5 including fresh protease inhibitors. Sample protein concentrations were determined using the BCA assay (ThermoScientific). For each pulldown, 3 mg of cell lysate was pre-incubated with either DMSO control, 20 μM UNC7145, or 20 μM UNC6934 (final concentration) for 1 hour with rotation at 4°C. For each sample, 25 μl of M270 Dynabeads (ThermoScientific) were prepared by washing three times in low salt wash buffer (10 mM Tris-HCl pH7.9, 100 mM NaCl, 0.1% NP-40), followed by incubation with 1 μM UNC7096 (biotinylated probe) for 1 hour at 4 °C. The unbound biotinylated compound was removed by 3 washes with low salt buffer. UNC7096 bound beads were then added to each sample followed by incubation 1 hour with rotation at 4oC. Beads were then washed 3 times with low-salt wash buffer followed by 2 washes with 50mM ammonium bicarbonate. On-bead digestion was performed by overnight incubation at 37°C with 2 μg of mass spectrometry grade trypsin (Promega). The following morning an additional 2 μg of trypsin was added to each sample and incubated at 37°C for 4-6 hours. The supernatant, containing digested peptides, was collected. Beads were then washed twice with water and supernatant pooled with digested peptides. Samples were then acidified with formic acid to a final concentration of 2% final concentration and flash frozen prior drying under vacuum.

### Label-free quantitative mass spectrometry data analysis

Raw MS/MS files were searched and quantified using Maxquant version 1.6.7.0 using the UP000005640 Uniprot human database (containing 20,605 protein entries, last modified November 5, 2019) with label-free quantification enabled and variable modifications oxidized methionine (+15.9949 Da) and deamidated asparagine (+0.9840) set. First search peptide tolerance and main search peptide tolerance were set at 30 and 6 ppm, respectively. For all other parameters default settings were used.

Differential enrichment analysis was performed using the DEP package (v1.8.0) in R (v3.5.1). Briefly, samples were filtered for proteins identified in 2 out of 3 replicates of at least one condition, normalized by variance stabilizing normalization and tested for differential enrichment relative to pulldowns competed with DMSO vehicle control.

### Fluorescence microscopy

For immunofluorescence, cells were fixed with 2% formaldehyde in 1x phosphate buffered saline (PBS) for 10 minutes at room temperature, followed by 3 washes in 1x PBS and permeabilization with 0.25% Triton X-100 in 1x PBS. Samples were then blocked with 1% bovine serum albumin (BSA; Sigma) in PBS-T (1x PBS and 0.1% Tween 20) for 1 hour at room temperature and incubated overnight with primary antibodies staining for NSD2 (Abcam Cat# ab75359, RRID:AB_1310816, 1:500) and fibrillarin (Abcam Cat# ab5821, RRID:AB_2105785, 1:1000). Cells were washed three times with 1x PBS, followed by staining with fluorescent anti-rabbit (Alexa Fluor^®^ 647 Conjugate; Cell Signalling Technologies; # 4414, RRID:AB_10693544, 1:1000) and anti-mouse (Alexa Fluor^®^ 488 Conjugate; Cell Signalling Technologies; #4408, RRID:AB_ 10694704, 1:1000) for 1 hour at room temperature. Unbound secondary antibodies were removed by three 1x PBS washes and DNA stained with Hoechst 33342 (Cell Signalling Technologies).

For microscopy of fluorescent fusion proteins, U2OS cells were seeded in 48-well plates 24 hours before transfection with 0.125 μg of NSD2-GFP and 0.125 μg of Fibrillarin-RFP using X-tremeGENE™ 360 following the manufacturer’s guidelines (Millipore Sigma). Cells were incubated an additional 24 hours before compound treatment or direct fixation. Cells were fixed with 2% formaldehyde in 1x PBS for 10 minutes at room temperature. DNA was then stained with Hoechst 33342 (Cell Signalling Technologies).

For confocal microscopy, images were acquired with Quorum Spinning Disk Confocal Microscope equipped with 405, 491, 561, and 642 nm lasers (Zeiss) and processed with Volocity software (Perkin Elmer) and ImageJ. For localization measurements of fluorescent fusion proteins, images were acquired with a EVOS™ FL Auto 2 Imaging System (Thermo Scientific™ Invitrogen™). Co-localization measurements were quantified using a custom CellProfiler (v3.1.9)^47^ analysis pipeline.

### Western blotting for global H3K36me2

Cells treated with compound for 72 hours before harvesting by centrifugation at 300 x g for 5 min. Cells were washed in 1x PBS and resuspended in lysis buffer containing 20 mM Tris-HCl (pH 8.0), 150 mM NaCl, 1 mM EDTA, 10 mM MgCl_2_, 0.5% TritonX-100, 30 μg/μl benzonase, and fresh protease inhibitors. After incubation for 3 min at room temperature, sodium dodecyl sulfate (SDS) was added to a final concentration of 1%. Samples were boiled in SDS loading buffer before western blotting using the NuPAGE electrophoresis and transfer system (Invitrogen) and near-infrared detection for H3K36me2 (Abcam Cat# ab9049, RRID:AB_1280939, 1:2000), Histone H3 (Abcam Cat# ab10799, RRID:AB_470239, 1:5000), alpha-tubulin (DSHB Cat# E7, RRID:AB_528499, 1:5000), and NSD2 (Abcam Cat# ab75359, RRID:AB_1310816), 1:1000). Immunoblots were imaged on a LiCor Odyssey CLx and quantified in Image Studio Lite v5.2.5 (Li-Cor Biosciences).

### Proliferation of KMS11 co-cultured with OP9 murine bone marrow stromal cells

KMS11 parental and TKO2, harbouring a knockout NSD2 expression from the t(4;14) allele, were transduced with GFP lentivirus and cells FACS sorted to obtain a homogenous GFP-expressing population. Cells were treated with compound at 5 μM for ten days before seeding on confluent monolayer of OP9 bone marrow stromal cells at density of 2000 cells per well of a 96 well plate. Proliferation was monitored by counting GFP-expressing cells over time using an IncuCyte live-cell imaging and analysis platform (Sartorius).

### 5-ethyl uridine (5-EU) incorporation assay

5-EU incorporation assays to measure changes in nucleolar transcription were performed as previously described^48^. To measure nucleolar transcription, cells were pre-treated with either 5 μM **UNC6934, UNC7145** or DMSO vehicle for 30 minutes and then fed 5-ethyl uridine (5-EU) for an additional 45 minutes. Cells were fixed with 2% paraformaldehyde (PFA) in PBS for 10 min at room temperature and then stained for nucleolin. Nascent RNA was labeled using the Click-iT RNA Alexa Fluor 594 imaging kit (Thermo Fisher Scientific) as per the supplier’s instructions. EVOS™ FL Auto 2 Imaging System (Thermo Scientific™ Invitrogen) and ribosomal RNA (rRNA) was defined by generating a mask of the 5-EU signal overlap with nucleolin. Image pre-processing, masking, and intensity measurements were done using a custom CellProfiler (v3.1.9)^47^ analysis pipeline

### RT-qPCR

Total RNA was first isolated using the Monarch Total RNA Miniprep (New England BioLabs) following the manufacturer’s protocol. Isolated RNA was reverse transcribed using the iScript cDNA Synthesis Kit (Bio-Rad). RT-qPCR was performed using a CFX384 Touch Real-Time PCR Detection System (Bio-Rad) and PowerUp SYBR Green Master Mix (Thermo Fisher, 100029284). Relative transcript levels were determined by the delta-delta Ct method and normalized to the housekeeping gene β2 microglobulin. Primer sequences are shown in Supplementary Table 3.

### Cell Fractionation

KMS11 cell pellets were washed in 1x PBS and resuspended in 200 μL of Hypotonic Buffer A (10 mM HEPES pH 7.5, 10 mM KCl, 1.5 mM MgCl_2_, 0.3 M Sucrose, 1 mM TCEP, and protease inhibitors) per 1 x 10^6^ cells. The cell suspension was incubated on ice for 15 minutes followed 1300 x g for 5 minutes. The supernatant was collected and cleared by centrifugation to produce the cytoplasmic (S1) fraction. The pellet was washed in Buffer A and resuspended in an equal volume of Buffer B-No NaCl (3 mM EDTA, 3 mM EGTA, 1 mM TCEP, and protease inhibitors) with/without 10 μM **UNC6934** or **UNC7145** negative control. Following incubation on ice for 30 minutes, the S2 fraction was centrifuged at 1300 x g for 5 minutes. The supernatant was again collected and pellet resuspended in an equal volume of Buffer B + NaCl (150 mM NaCl, 3 mM EDTA, 3 mM EGTA, 1 mM TCEP, and protease inhibitors) with/without 10 μM **UNC6934** or **UNC7145** negative control. Following incubation on ice for 10 minutes, samples were centrifuged at 1300 x g for 5 minutes with the supernatent yielding the salt-extracted nucleoplasmic (S3) fraction. The chromatin-bound or pellet fraction (P) was then resuspended in an equal volume of Histone buffer (20 mM Tris HCl pH 7.5, 150 mM NaCl, 1 mM EDTA, 10 mM MgCl2, 1% Triton X-100, 1 mM TCEP, 30 μg.μL^-1^ benzonase, and protease inhibitors). Samples were boiled in SDS loading buffer before western blotting with antibodies to NSD2 (Abcam Cat# ab75359, RRID:AB_1310816, 1:1000) using the NuPAGE electrophoresis and transfer system (Invitrogen). Immunoblots were imaged on a LiCor Odyssey CLx and quantified in Image Studio Lite v5.2.5 (Li-Cor Biosciences).

#### Chemistry

##### General Chemistry Procedures

Reactions were carried out using conventional glassware. All reagents and solvents were used as received unless otherwise stated. Reagents were of 95% purity or greater, and solvents were reagent grade unless otherwise stated. Any anhydrous solvents used were purchased as “anhydrous” grade and used without further drying. “Room” or ambient temperature varied between 20-25°C. Analytical thin layer chromatography (TLC) was carried out using glass plates pre-coated with silica gel (Merck) impregnated with fluorescent indicator (254 nm). TLC plates were visualized by illumination with a 254 nm UV lamp. Analytical LCMS data for all compounds were acquired using an Agilent 1260 Infinity II system with the UV detector set to 254 nm. Samples were injected (<10 μL) onto an Agilent ZORBAX Eclipse Plus C18, 600 Bar, 4.6 x 50 mm, 1.8 μM column at 25.0°C. Mobile phases A (H_2_O + 0.1% acetic acid), B (MeOH + 0.1% acetic acid), and C (99% MeCN + 1% H_2_O + 0.1% acetic acid) were used with a linear gradient from 10% to 100% B or C in 5.0 min, followed by a flush at 100% B or C for another 2 minutes with a flow rate of 1.0 mL/min. Mass spectra (MS) data were acquired in positive ion mode using an Agilent InfinityLab LC/MSD single quadrupole mass spectrometer with an electrospray ionization (ESI) source. Normal phase column chromatography was performed with a Teledyne Isco CombiFlash^®^R_f_ 200 using RediSep^®^R_f_ SILICA columns with the UV detector set to 254 nm and 280 nm. Reverse phase column chromatography was performed with a Teledyne Isco CombiFlash^®^R_f_ 200 using C18 RediSep^®^R_f_ Gold columns with the UV detector set to 220 nm and 254 nm. Analytical LCMS (at 254 nm) was used to establish the purity of targeted compounds. All compounds that were evaluated in biochemical and biophysical assays had >95% purity as determined by LCMS (spectra provided in supplementary information).

#### Nuclear Magnetic Resonance Spectroscopy (NMR)

^1^H and ^13^C NMR spectra were obtained on a Varian 400MR at 400 MHz and 101 MHz respectively. Chemical shifts are reported in ppm and coupling constants are reported in Hz with CDCl_3_ referenced at 7.26 (^1^H) and 77.1 ppm (^13^C), DMSO-*d*_6_ referenced at 2.50 (^1^H) and 39.5 ppm (^13^C), acetone-*d*_6_ referenced at 2.05 (^1^H) and 29.8 ppm (^13^C), and MeOD-*d*_4_ referenced at 3.31 (^1^H) and 49.0 ppm (^13^C). All compounds that were evaluated in biochemical and biophysical assays had >95% purity as determined by ^1^H NMR (spectra provided in supplementary information).

#### High-Resolution Mass Spectrometry

Samples were prepared by dilution with 600 μL of acetonitrile. Samples were further diluted using 100 μL of the compound solution in 900 μL of acetonitrile, regardless of the original dilution solvent.

Samples were analyzed with a ThermoFisher Q Exactive HF-X (ThermoFisher, Bremen, Germany) mass spectrometer coupled with a Waters Acquity H-class liquid chromatograph system. Samples were introduced via a heated electrospray source (HESI) at a flow rate of 0.6 mL/min. Electrospray source conditions were set as: spray voltage 3.0 kV, sheath gas (nitrogen) 60 arb, auxillary gas (nitrogen) 20 arb, sweep gas (nitrogen) 0 arb, nebulizer temperature 375 degrees C, capillary temperature 380 degrees C, RF funnel 45 V. The mass range was set to 150-2000 m/z. All measurements were recorded at a resolution setting of 120,000.

Separations were conducted on a Waters Acquity UPLC BEH C18 column (2.1 x 50 mm, 1.7 um particle size). LC conditions were set at 95% water with 0.1% formic acid (A) ramped linearly over 5.0 mins to 100% acetonitrile with 0.1% formic acid (B) and held until 6.0 mins. At 7.0 mins the gradient was switched back to 95% A and allowed to re-equilibrate until 9.0 mins. The injection volume for all samples was 3 μL.

Xcalibur (ThermoFisher, Breman, Germany) was used to analyze the data. Solutions were analyzed at 0.1 mg/mL or less based on responsiveness to the ESI mechanism. Molecular formula assignments were determined with Molecular Formula Calculator (v 1.2.3). All observed species were singly charged, as verified by unit *m/z* separation between mass spectral peaks corresponding to the ^12^C and ^13^C^12^C_c-1_ isotope for each elemental composition.

## Abbreviations used

DCM: Dichloromethane
DIPEA: *N,N*-Diisopropylethylamine
DMAP: 4-(Dimethylamino)pyridine
DMF: *N,N*-Dimethylformamide
DMSO: Dimethylsulfoxide
EDC: *N*-(3-Dimethylaminopropyl)-*N’*-ethylcarbodiimide hydrochloride
HOAt: 1-Hydroxy-7-azabenzotriazole
LCMS: Liquid Chromatography-Mass Spectrometry
NMR: Nuclear Magnetic Resonance
TCFH: Chloro-*N,N,N’,N’*-tetramethylformamidinium hexafluorophosphate
TFA: 2,2,2-Trifluoroacetic acid
THF: Tetrahydrofuran

## Synthetic Schemes

**Supplemental Scheme 1:**
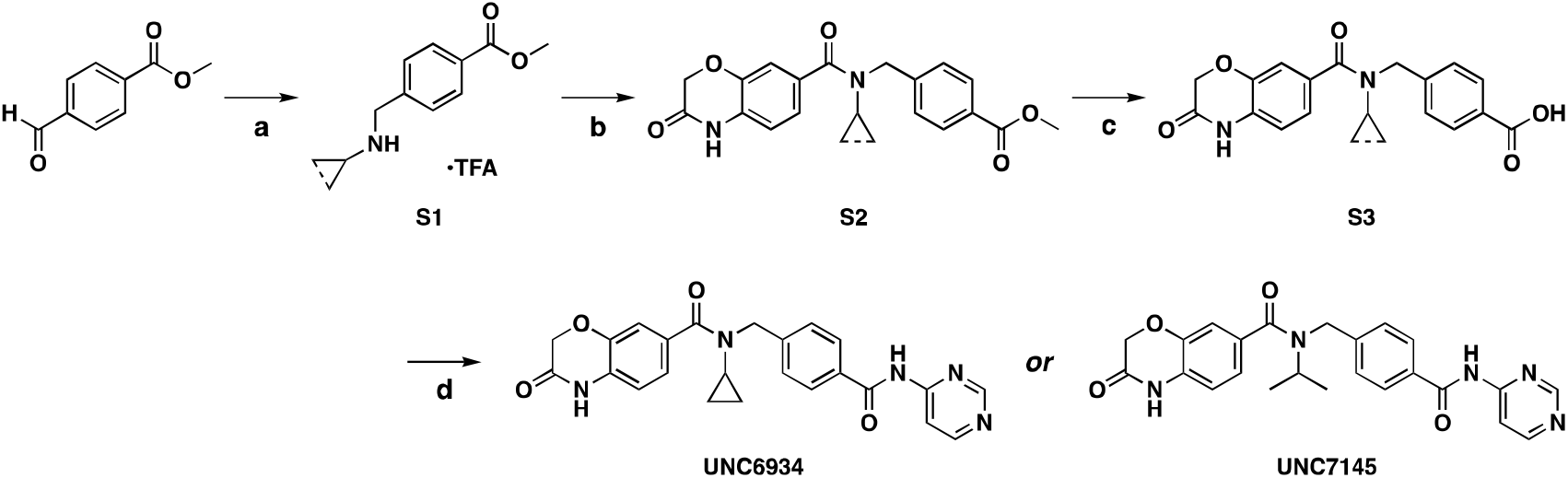
Synthesis of probe compounds. Reagents and conditions: a) i. (cyclo/iso)propylamine, MeOH ii. NaBH_4_; b) 3-oxo-3,4-dihydro-2*H*-benzo[*b*][1,4]oxazine-7-carboxylic acid, EDC, HOAt, triethylamine, MeCN; c) LiOH•H_2_O; d) 4-aminopyrimidine, 1-methylimidazole, TCFH^49^. Note: Cyclopropyl-containing intermediates are designated with a ‘*P*’ (probe) and isopropyl-containing intermediates are designated with an ‘*N*’ (negative control).

**Supplemental Scheme 2:**
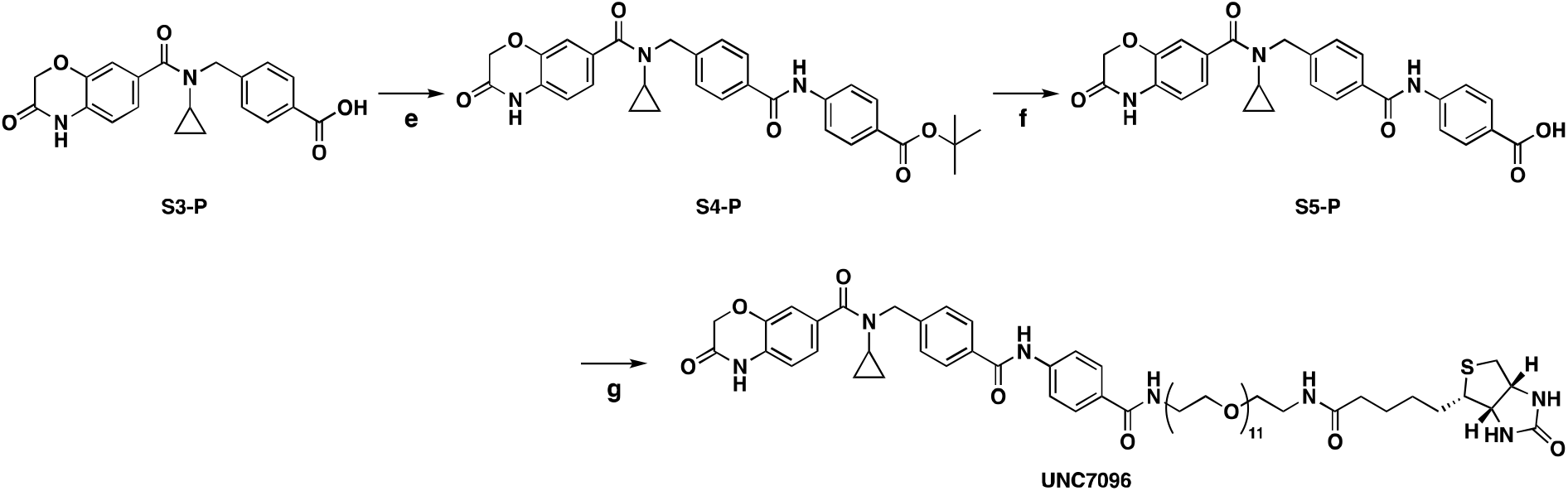
Synthesis of biotinylated tool compound UNC7096. Reagents and conditions: e) *tert*-butyl 4-aminobenzoate, EDC, DMAP, DMF; f) TFA, DCM; g) Biotin-PEG11-amine, EDC, HOAt, DIPEA, DMF.

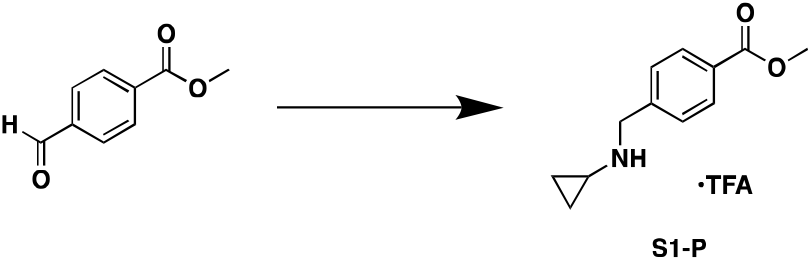

To a 50 mL flask equipped with a stir bar was added methyl 4-formylbenzoate (1.0 g, 1 Eq, 6.1 mmol) and methanol (10 mL), followed by cyclopropylamine (0.35 g, 0.43 mL, 1 Eq, 6.1 mmol). The flask was capped and stirred at room temperature overnight. The next day, the flask was cooled in an ice water bath and sodium borohydride (0.46 g, 2 Eq, 12 mmol) was added portionwise. Borohydride addition was accompanied by effervescence and heating of the solution. After 4 hours, at which time the reaction had come to room temperature, the reaction was quenched by addition of saturated sodium bicarbonate and extracted three times with ethyl acetate. The combined organic layers were washed once more with saturated sodium bicarbonate, once with brine, then dried over sodium sulfate and concentrated to an oil. Normal phase chromatography over silica (0-100% ethyl acetate in hexanes) provided the free base as a colorless free-flowing oil. The oil was dissolved in 25 mL of diethyl ether and cooled in an ice water bath, and trifluoroacetic acid (1.2 g, 0.80 mL, 1.7 Eq, 10 mmol) was added dropwise. A voluminous white solid formed, which was filtered and washed rigorously with diethyl ether to provide **S1-P** (1.586 g, 4.976 mmol, 82%) as a fluffy white powder.

^1^H NMR (400 MHz, Methanol-*d*_4_) δ 8.09 (d, *J* = 8.5 Hz, 2H), 7.62 (d, *J* = 8.4 Hz, 2H), 4.39 (s, 2H), 3.92 (s, 3H), 2.83 – 2.75 (m, 1H), 0.96 – 0.85 (m, 4H).

^13^C NMR (101 MHz, Methanol-*d*_4_) δ 167.74, 137.52, 132.47, 131.24, 131.19, 52.84, 52.36, 31.25, 4.27.

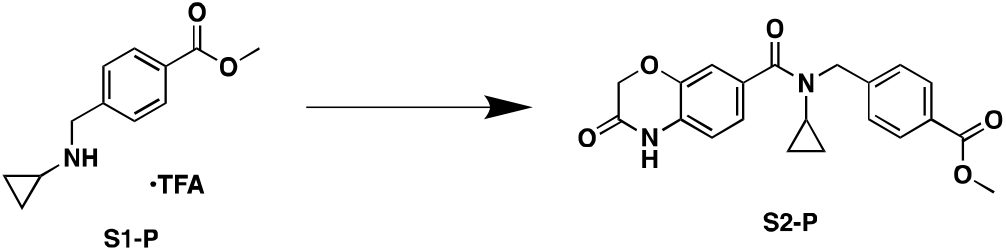

To a flask was added 3-oxo-3,4-dihydro-2*H*-benzo[*b*][1,4]oxazine-7-carboxylic acid (200 mg, 1 Eq, 1.04 mmol), HOAt (211 mg, 1.5 Eq, 1.55 mmol), EDC (298 mg, 1.5 Eq, 1.55 mmol), and acetonitrile (5 mL). The mixture was stirred for 15 minutes at room temperature to preactivate the acid. To the flask was then added **S1-P** (397 mg, 1.2 Eq, 1.24 mmol) and triethylamine (314 mg, 0.43 mL, 3 Eq, 3.11 mmol), and the reaction stirred at room temperature overnight. The next day, the reaction was quenched with water and extracted 3 times with ethyl acetate. The combined organic fractions were washed once with 0.5 M citric acid, once with water, once with saturated sodium bicarbonate, and once with brine, then dried over sodium sulfate, filtered, and concentrated to a white solid. Normal phase chromatography over silica (0-100% ethyl acetate in DCM) provided **S2-P** (292.7 mg, 769.5 μmol, 74.3%) as a white solid.

^1^H NMR (400 MHz, Chloroform-*d*) δ 9.43 (s, 1H), 8.02 (d, *J* = 8.3 Hz, 2H), 7.39 (d, *J* = 7.9 Hz, 2H), 7.19 - 7.12 (m, 2H), 6.85 (d, *J* = 7.8 Hz, 1H), 4.77 (s, 2H), 4.62 (s, 2H), 3.91 (s, 3H), 2.62 (tt, *J* = 6.9, 4.0 Hz, 1H), 0.67 - 0.55 (m, 2H), 0.55 - 0.45 (m, 2H).

^13^C NMR (101 MHz, Chloroform-*d*) δ 167.00, 166.05, 143.15, 132.88, 130.15, 129.83, 129.45, 127.96, 127.69, 122.40, 116.35, 115.76, 67.27, 52.28, 27.02, 10.15. 1 aromatic and 1 aliphatic C not observed.

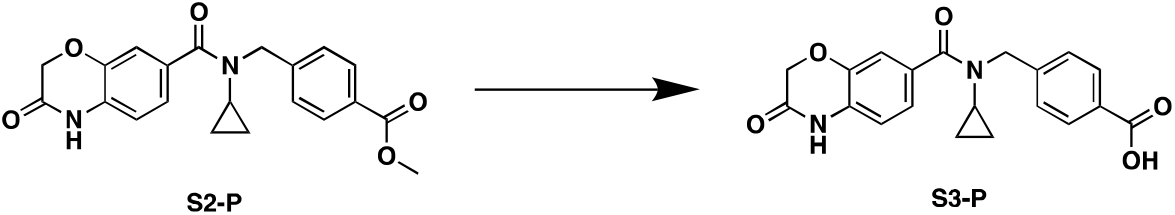

To a round-bottom flask was added **S2-P** (904 mg, 1 Eq, 2.38 mmol), THF (20 mL), and lithium hydroxide hydrate (10% in water, 5 mL, 5 Eq, 11.9 mmol). The reaction was stirred at room temperature for 24 hours, then extracted 5 times with 20 mL portions of diethyl ether. The aqueous layer was then acidified to pH 2 with 1 M KHSO_4_, and the precipitate produced was filtered off, washed with cold water, and dried to provide **S3-P** (771.1 mg, 2.105 mmol, 88.6%) as a white solid.

^1^H NMR (400 MHz, DMSO-*d*_6_) δ 10.87 (s, 1H), 7.93 (d, *J* = 8.2 Hz, 2H), 7.41 (d, *J* = 8.0 Hz, 2H), 7.18 (dd, *J* = 8.1, 1.8 Hz, 1H), 7.14 (d, *J* = 1.7 Hz, 1H), 6.92 (d, *J* = 8.1 Hz, 1H), 4.70 (s, 2H), 4.61 (s, 2H), 2.84 – 2.73 (m, 1H), 0.59 – 0.49 (m, 2H), 0.49 – 0.38 (m, 2H).

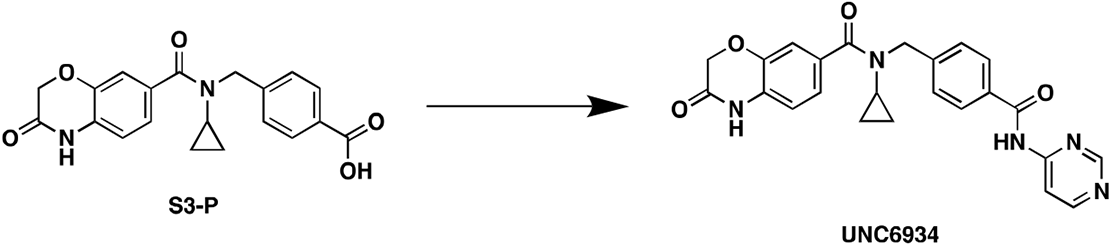

To a 2-dram vial with a stirbar was added pyrimidin-4-amine (50.6 mg, 1.3 Eq, 532 μmol), **S3-P** (150 mg, 1 Eq, 409 μmol), 1-methylimidazole (118 mg, 114 μL, 3.5 Eq, 1.43 mmol), THF (1.5 mL), and finally TCFH (149 mg, 1.3 Eq, 532 μmol). The reaction was left to stir at room temperature for 24 hours, then quenched with 25 mL of distilled water and extracted 7 times with 25 mL portions of ethyl acetate. The combined organic layers were extracted once with 15 mL of saturated sodium bicarbonate (for recovery of the starting acid), then washed twice with water, once with saturated sodium bicarbonate, and once with brine, and finally dried over anhydrous sodium sulfate and concentrated to a yellow oil. The oil was dry-loaded on Celite and purified by normal phase chromatography over silica (0-10% methanol in DCM) to yield **UNC6934** (106 mg, 239 μmol, 58.4%) as an off-white solid.

^1^H NMR (400 MHz, DMSO-*d*_6_) δ 11.21 (s, 1H), 10.87 (s, 1H), 8.95 (d, *J* = 1.4 Hz, 1H), 8.72 (d, *J* = 5.8 Hz, 1H), 8.21 (dd, *J* = 5.8, 1.3 Hz, 1H), 8.03 (d, *J* = 8.3 Hz, 2H), 7.45 (d, *J* = 8.0 Hz, 2H), 7.19 (dd, *J* = 8.0, 1.8 Hz, 1H), 7.15 (d, *J* = 1.7 Hz, 1H), 6.93 (d, *J* = 8.1 Hz, 1H), 4.72 (s, 2H), 4.62 (s, 2H), 2.88 – 2.77 (m, 1H), 0.61 – 0.50 (m, 2H), 0.50 – 0.41 (m, 2H).

^13^C NMR (100 MHz, DMSO-*d*_6_) δ 166.90, 164.85, 158.30, 158.26, 158.17, 143.21, 142.48, 132.03, 131.73, 128.63, 128.40, 127.16, 121.90, 115.39, 115.14, 110.65, 66.73, 9.54. 1 aromatic C and 2 aliphatic C not observed.

LCMS (ESI, +ve mode) expected *m/z* for C_24_H_22_N_5_O_4_ [M+H] 444.17, found 444.15

HRMS (ESI, +ve mode) expected *m/z* for C_24_H_22_N_5_O_4_ [M+H] 444.16663, found 444.16638

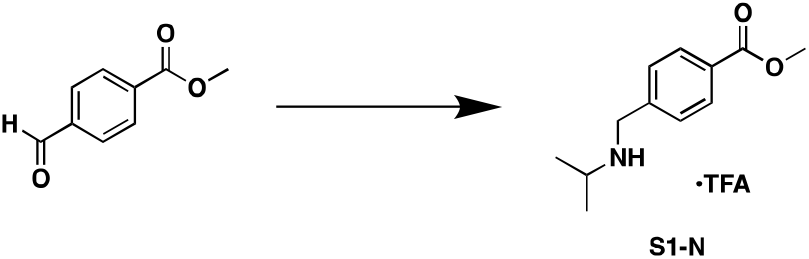

To a 50 mL flask equipped with a stir bar was added methyl 4-formylbenzoate (0.50 g, 1 Eq, 3.0 mmol) and methanol (5 mL), followed by isopropylamine (0.18 g, 0.26 mL, 1 Eq, 3.0 mmol). The flask was capped and stirred at room temperature overnight. The next day, the flask was cooled in an ice water bath and sodium borohydride (0.23 g, 2 Eq, 6.1 mmol) was added portionwise. Borohydride addition was accompanied by effervescence and heating of the solution. After 4 hours, at which time the reaction had come to room temperature, the reaction was quenched by addition of saturated sodium bicarbonate, and extracted three times with ethyl acetate. The combined organic layers were washed once more with saturated sodium bicarbonate, once with brine, then dried over sodium sulfate and concentrated to an oil. Normal phase chromatography over silica (0-100% ethyl acetate in hexanes) provided the free base as a colorless free-flowing oil. The oil was dissolved in 25 mL of diethyl ether and cooled in an ice water bath, and trifluoroacetic acid (0.52 g, 0.35 mL, 1.5 Eq, 4.6 mmol) was added dropwise. A voluminous white solid formed, which was filtered and washed rigorously with diethyl ether to provide **S1-N** (773.4 mg, 2.407 mmol, 79%) as a fluffy white powder.

^1^H NMR (400 MHz, Methanol-*d*_4_) δ 8.10 (d, *J* = 8.4 Hz, 2H), 7.62 (d, *J* = 8.6 Hz, 2H), 4.29 (s, 2H), 3.92 (s, 3H), 3.53 - 3.42 (m, 1H), 1.41 (d, *J* = 6.6 Hz, 6H).

^13^C NMR (100 MHz, Methanol-*d*_4_) δ 167.75, 137.93, 132.38, 131.23, 131.01, 52.83, 52.23, 49.15, 19.21.

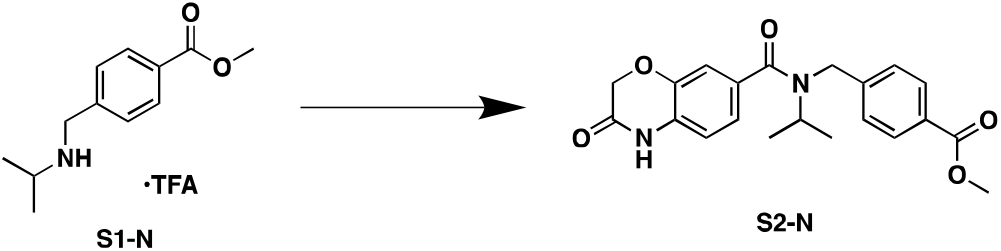

To a scintillation vial was added 3-oxo-3,4-dihydro-2*H*-benzo[*b*][1,4]oxazine-7-carboxylic acid (50 mg, 1 Eq, 0.26 mmol), EDC (87 mg, 1.75 Eq, 0.45 mmol), HOAt (70 mg, 2 Eq, 0.52 mmol), and acetonitrile (2.0 mL). The reaction was left to stir at room temperature for 30 minutes. To the vial was then added **S1-N** (0.10 g, 1.2 Eq, 0.31 mmol), followed by triethylamine (0.10 g, 0.14 mL, 4 Eq, 1.0 mmol), and the reaction was left to stir overnight. The next day, the volatiles were removed under reduced pressure and the residue was partitioned between water and ethyl acetate. The layers were separated and the aqueous layer was extracted twice more with ethyl acetate. The combined organic layers were washed once with 0.5 M citric acid, once with water, once with saturated sodium bicarbonate, and once with brine, then dried over sodium sulfate and concentrated to an off-white residue. Normal phase chromatography over silica gel (0-100% ethyl acetate in DCM) provided **S2-N** (90.1 mg, 236 μmol, 91%) as a white solid.

^1^H NMR (400 MHz, Chloroform-*d*) δ 9.58 (s, 1H), 8.01 - 7.94 (m, 2H), 7.36 (br s, 2H), 7.01 (d, *J* = 15.7 Hz, 2H), 6.84 (br s, 1H), 4.65 (s, 2H), 4.61 (s, 2H), 4.21 (s, 1H), 3.89 (s, 3H), 1.13 (d, *J* = 6.6 Hz, 6H).

^13^C NMR (100 MHz, Chloroform-*d*) δ 171.38, 167.02, 165.94, 144.52, 143.57, 132.68, 129.94, 128.97, 127.48, 126.93, 121.00, 116.31, 115.19, 67.21, 52.21, 21.43, 14.29. 1 aliphatic C not observed.

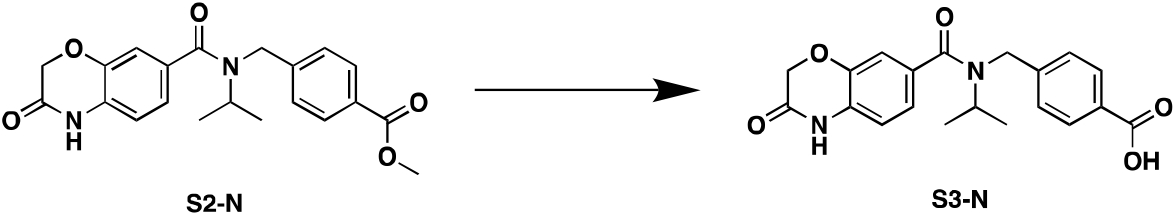

To a round-bottom flask was added **S2-N** (405 mg, 1 Eq, 1.06 mmol), 1,4-dioxane (6 mL), and lithium hydroxide hydrate (15% in water, 1.5 mL, 5 Eq, 5.30 mmol). The reaction was stirred at room temperature for 24 hours, then extracted 5 times with 20 mL portions of diethyl ether. The aqueous layer was then acidified to pH 2 with 1 M KHSO_4_, and the precipitate produced was filtered off, washed with cold water, and dried to provide **S3-N** (319.8 mg, 868.1 μmol, 82.0%) as a white solid.

^1^H NMR (400 MHz, DMSO-*d*_6_) δ 10.85 (s, 1H), 7.89 (d, *J* = 8.2 Hz, 2H), 7.41 (d, *J* = 7.7 Hz, 2H), 7.03 (s, 2H), 6.99 - 6.89 (m, 1H), 4.61 (s, 4H), 4.08 (s, 1H), 1.08 (d, *J* = 6.6 Hz, 6H).

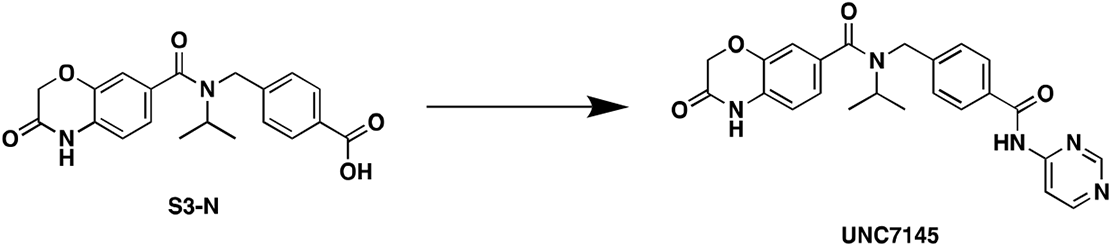

To a 2-dram vial with a stirbar was added pyrimidin-4-amine (33.6 mg, 1.3 Eq, 353 μmol), **S3-N** (100 mg, 1 Eq, 271 μmol), 1-methylimidazole (78.0 mg, 75.7 μL, 3.5 Eq, 950 μmol), acetonitrile (1.0 mL), and finally TCFH (99.0 mg, 1.3 Eq, 353 μmol). The reaction was heated to 40 °C for 16 hours, then quenched with 25 mL of distilled water and extracted 6 times with 25 mL portions of ethyl acetate. The combined organic layers were extracted once with 15 mL of saturated sodium bicarbonate (for recovery of the starting acid), then washed twice with water, once with saturated sodium bicarbonate, and once with brine, and finally dried over anhydrous sodium sulfate and concentrated to a yellow oil. The oil was dry-loaded on Celite and purified by normal phase chromatography over silica (0-10% methanol in DCM) to yield **UNC7145** (52 mg, 0.12 mmol, 43%) as an off-white solid.

^1^H NMR (400 MHz, DMSO-*d*_6_) δ 11.18 (s, 1H), 10.86 (s, 1H), 8.95 (d, *J* = 1.3 Hz, 1H), 8.71 (d, *J* = 5.8 Hz, 1H), 8.21 (dd, *J* = 5.8, 1.3 Hz, 1H), 7.98 (d, *J* = 8.4 Hz, 2H), 7.51 – 7.39 (m, 2H), 7.10 – 6.99 (m, 2H), 6.96 (s, 1H), 4.69 – 4.54 (m, 4H), 4.16 – 4.03 (m, 1H), 1.10 (d, *J* = 6.6 Hz, 6H).

^13^C NMR (101 MHz, DMSO-*d*_6_) δ 170.23, 166.97, 164.78, 158.29, 158.27, 158.21, 144.61, 143.00, 131.76, 131.58, 128.36, 128.16, 126.58, 120.54, 115.76, 114.27, 110.67, 66.76, 50.47, 42.82, 20.74.

LCMS (ESI, +ve mode) expected *m/z* for C_24_H_24_N_5_O_4_ [M+H] 446.18, found 446.15

HRMS (ESI, +ve mode) expected *m/z* for C_24_H_24_N_5_O_4_ [M+H] 446.18228, found 446.18216

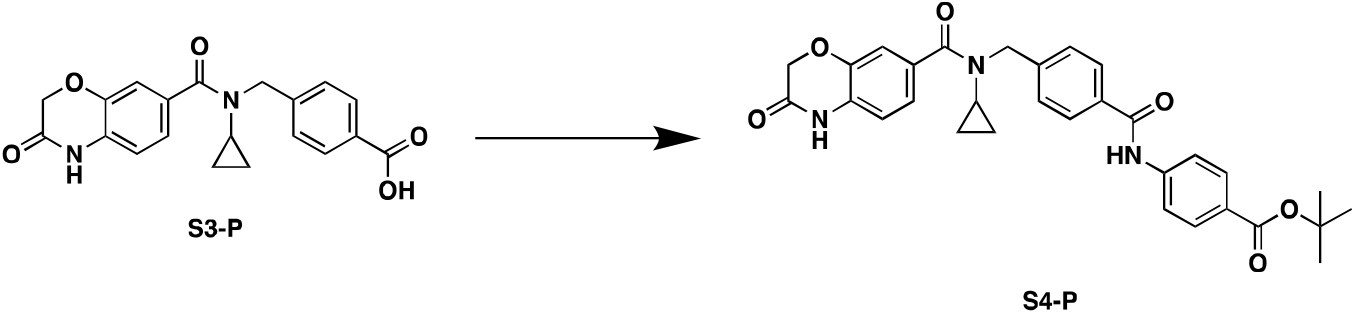

To a scintillation vial was added **S3-P** (18 mg, 1 Eq, 50 μmol), *tert-butyl* 4-aminobenzoate (19 mg, 2 Eq, 0.10 mmol), EDC (19 mg, 2 Eq, 0.10 mmol), DMAP (12 mg, 2 Eq, 0.10 mmol), and DMF (0.2 mL). The reaction was heated to 50 °C and stirred overnight. The next day, the reaction was partitioned between water and ethyl acetate. The layers were separated, and the aqueous layer was extracted twice more with ethyl acetate. The combined organic layers were washed twice with water, once with saturated sodium bicarbonate, and once with brine, then dried over sodium sulfate and concentrated to an off-white residue. Purification by normal phase chromatography over silica (0-100% ethyl acetate in DCM) provided **S4-P** (20 mg, 37 μmol, 74%) as a white solid.

^1^H NMR (400 MHz, Chloroform-*d*) δ 9.25 (s, 1H), 8.83 (s, 1H), 7.94 (d, *J* = 8.7 Hz, 2H), 7.84 (d, *J* = 8.3 Hz, 2H), 7.77 (d, *J* = 8.8 Hz, 2H), 7.31 (d, *J* = 7.9 Hz, 2H), 7.09 (s, 1H), 7.07 (d, *J* = 8.1 Hz, 1H), 6.79 (d, *J* = 8.0 Hz, 1H), 4.75 (s, 2H), 4.59 (s, 2H), 2.63 (tt, *J* = 7.0, 4.0 Hz, 1H), 1.59 (s, 9H), 0.66 - 0.56 (m, 2H), 0.56 - 0.44 (m, 2H).

^13^C NMR (100 MHz, Chloroform-*d*) δ 165.95, 165.68, 165.60, 143.15, 142.29, 141.99, 134.02, 132.57, 130.65, 127.95, 127.84, 127.64, 122.21, 119.43, 116.13, 115.81, 81.09, 67.29, 28.37. 2 aromatic C and 3 aliphatic C not observed.

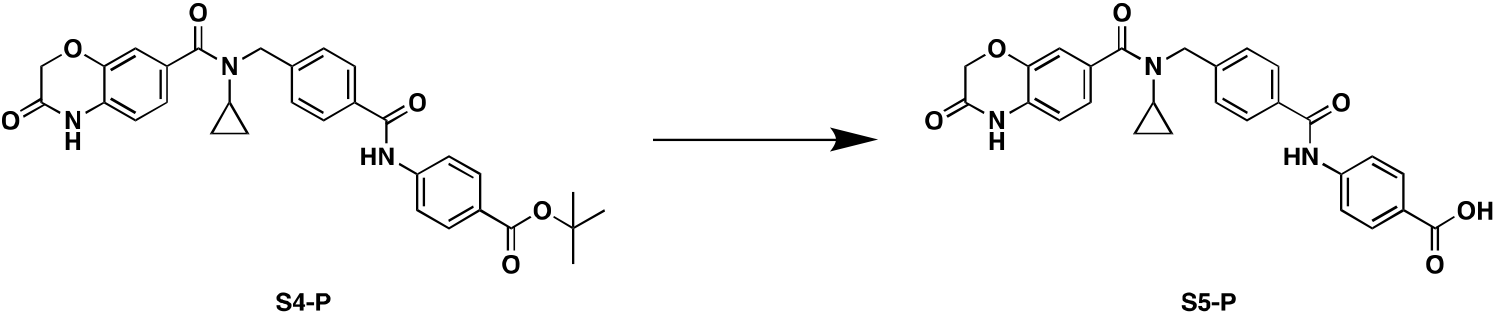

To a vial containing **S4-P** (155 mg, 1 Eq, 286 μmol) was added DCM (1.3 mL) and TFA (489 mg, 331 μL, 15 Eq, 4.29 mmol), which were stirred at room temperature overnight. The next day, the volatiles were removed *in vacuo* to provide **S5-P** (144 mg, 297 μmol, quantitative) as a tan solid.

^1^H NMR (400 MHz, DMSO-*d*_6_) δ 10.87 (s, 1H), 10.52 (s, 1H), 7.98 - 7.89 (m, 6H), 7.47 (d, *J* = 8.1 Hz, 2H), 7.18 (d, *J* = 8.3 Hz, 1H), 7.15 (d, *J* = 1.8 Hz, 1H), 6.93 (d, *J* = 8.1 Hz, 1H), 4.72 (s, 2H), 4.62 (s, 2H), 2.84 - 2.75 (m, 1H), 0.59 - 0.51 (m, 2H), 0.51 - 0.43 (m, 2H).

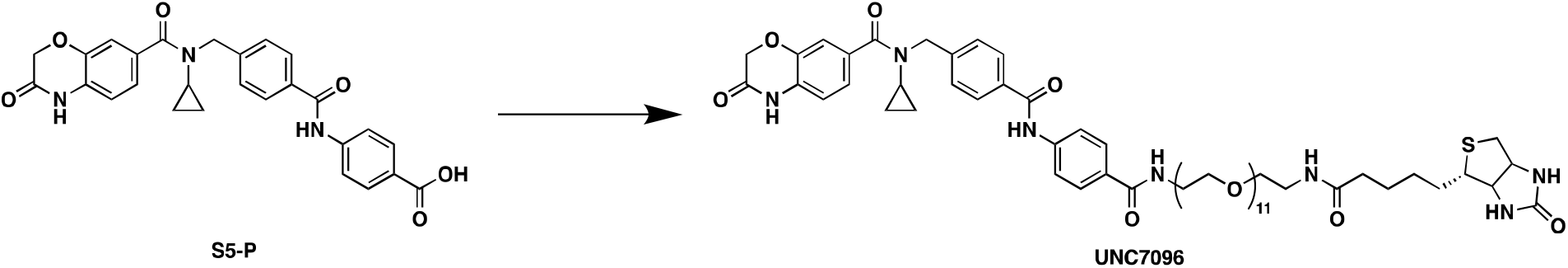

To a scintillation vial was added **S5-P** (3.1 mg, 1 Eq, 6.5 μmol), EDC (2.5 mg, 2 Eq, 13 μmol), HOAt (1.8 mg, 2 Eq, 13 μmol), and DMF (0.2 mL). The reaction was left to stir at room temperature for 30 minutes. To the vial was then added Biotin-PEG11-amine (5.0 mg, 1 Eq, 6.5 μmol) dissolved in DMF (0.2 mL), followed by DIPEA (2.5 mg, 3.4 μL, 3 Eq, 19 μmol), and the reaction was left to stir overnight. The next day, the reaction was diluted with distilled water and purified directly by reverse phase chromatography (10-100% methanol in water + 0.1% TFA) and lyophilized to provide **UNC7096** (6.51 mg, 5.26 μmol, 81%) as a white hygroscopic solid.

^1^H NMR (400 MHz, Methanol-*d*_4_) δ 7.97 (d, *J* = 8.3 Hz, 2H), 7.90 – 7.83 (m, 4H), 7.51 (d, *J* = 7.9 Hz, 2H), 7.20 (d, *J* = 8.6 Hz, 1H), 7.18 – 7.16 (m, 1H), 6.97 (d, *J* = 8.1 Hz, 1H), 4.83 (s, 2H), 4.63 (s, 2H), 4.48 (ddd, *J* = 7.9, 5.0, 0.9 Hz, 1H), 4.29 (dd, *J* = 7.9, 4.5 Hz, 1H), 3.71 – 3.55 (m, 40H), 3.53 (t, *J* = 5.4 Hz, 2H), 3.37 – 3.33 (m, 2H), 3.19 (dt, *J* = 9.8, 5.3 Hz, 1H), 2.92 (dd, *J* = 12.8, 5.0 Hz, 1H), 2.86 – 2.80 (m, 1H), 2.70 (d, *J* = 12.8 Hz, 1H), 2.21 (t, *J* = 7.4 Hz, 2H), 1.78 – 1.53 (m, 5H), 1.43 (p, *J* = 7.6 Hz, 2H), 0.66 (d, *J* = 6.5 Hz, 2H), 0.57 (s, 2H). PEG group underintegrates slightly.

LCMS (ESI, +ve mode) expected m/z for C_61_H_88_N_7_O_18_S+ [M+H] 1238.59, found 1238.40

## Statistics

For imaging experiments, Pearson correlation coefficients (PCC) were calculated using built-in functions in CellProfiler (v3.1.9). Welch t-tests to compare median PCC values of independent biological replicates were calculated with R (v3.5.1). For chemical proteomics experiments, adjusted p-values and enrichment scores were calculated using DEP package (v1.8.0) for proteomic analysis in R (v3.5.1).

## Data availability

The mass spectrometry proteomics data have been deposited to the ProteomeXchange Consortium via the PRIDE^50^ partner repository with the dataset identifier PXD017641.

The structure of NSD2-PWWP1 in complex with MRT866 and UNC06934 were deposited to the Protein Data Bank with accession number 7LMT and 6XCG respectively.

