## Supplementary information for "Pharmacological targeting of a PWWP domain demonstrates cooperative control of NSD2 localization"

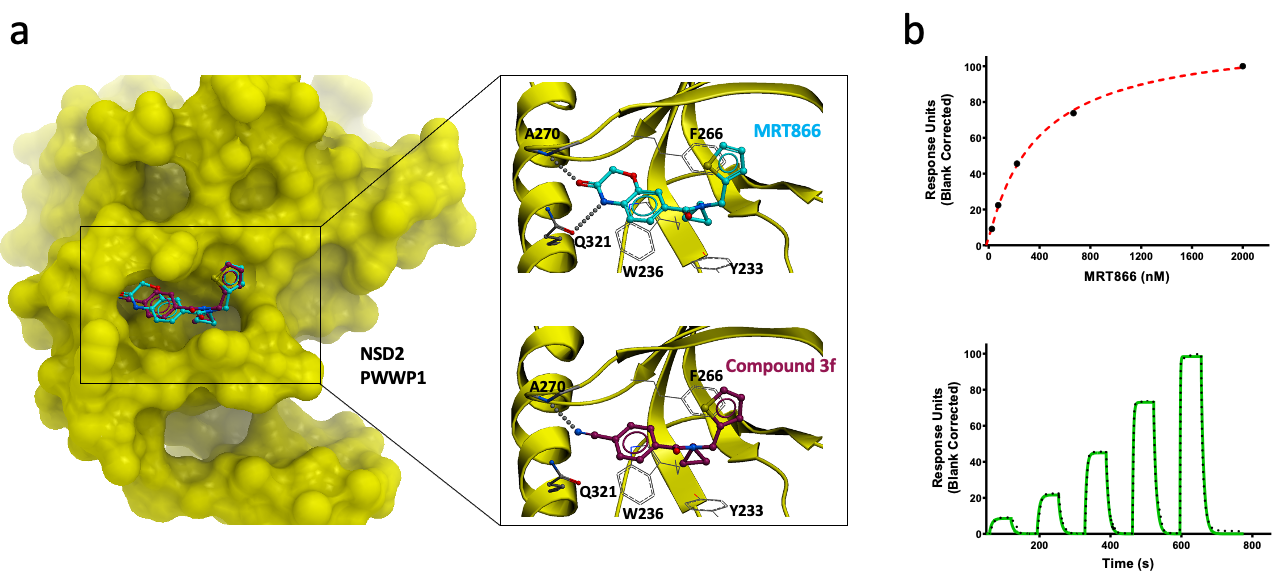

**Supplementary Figure 1: MRT866 binds NSD2-PWWP1**. a) Crystal structure of commercial compound **MRT866** bound to NSD2-PWWP1 (PDB 7LMT) and comparison with the structure of compound **3f** bound to the same site (PDB 6UE6). Aminoacids forming hydrogen-bonds (A270, Q321) and the aromatic cage (Y233,W236,F266) are shown. b) SPR analysis. Top: steady-state response (black circles) with the steady-state 1:1 binding model fitting (red dashed line). Bottom: representative sensorgram (green) with the kinetic fit (black dots). K_d_ value of 349 ± 19 nM was generated from kinetic fitting with the 1:1 binding model and averaged from triplicate.

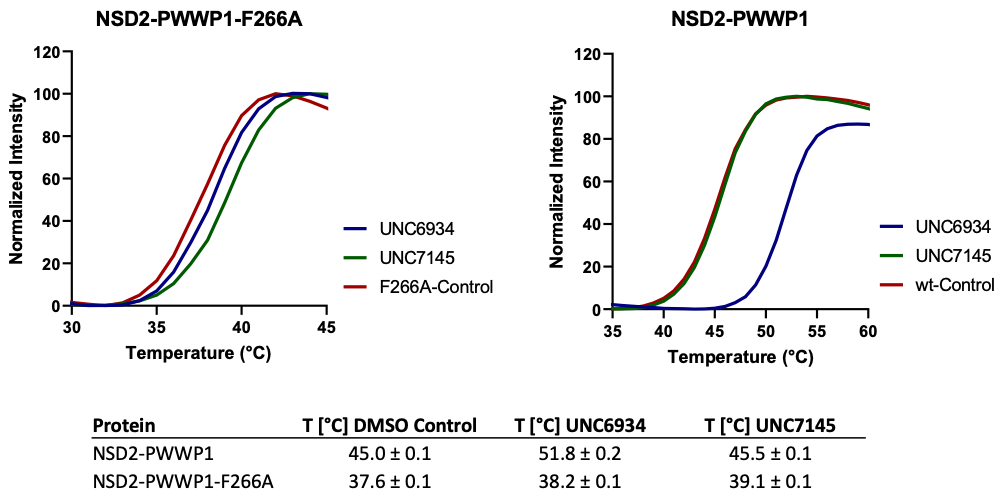

**Supplementary Figure 2: Binding of UNC6934 and UNC7145 to wt-NSD2-PWWP1 and its F266A mutant.** The binding of both compounds to NSD2-PWWP1 F266A mutant in parallel to the wild-type (wt) was assessed by DSF in quadruplicate at 100 µM of each compound. Assays were performed in 100 mM HEPES, 150 mM NaCl, pH 7.5, as previously described (https://doi.org/10.1038/nprot.2007.321). T_m_ (^o^C) values are presented as mean ± SD in the above table. High fluorescence background and weak transitions was observed in the presence of F266A mutant protein. The T_m_ of the F266A mutant (37.6 ± 0.1 ^o^C) was much lower than the wt-NSD2-PWWP1 (45 ± 0.1 ^o^C).

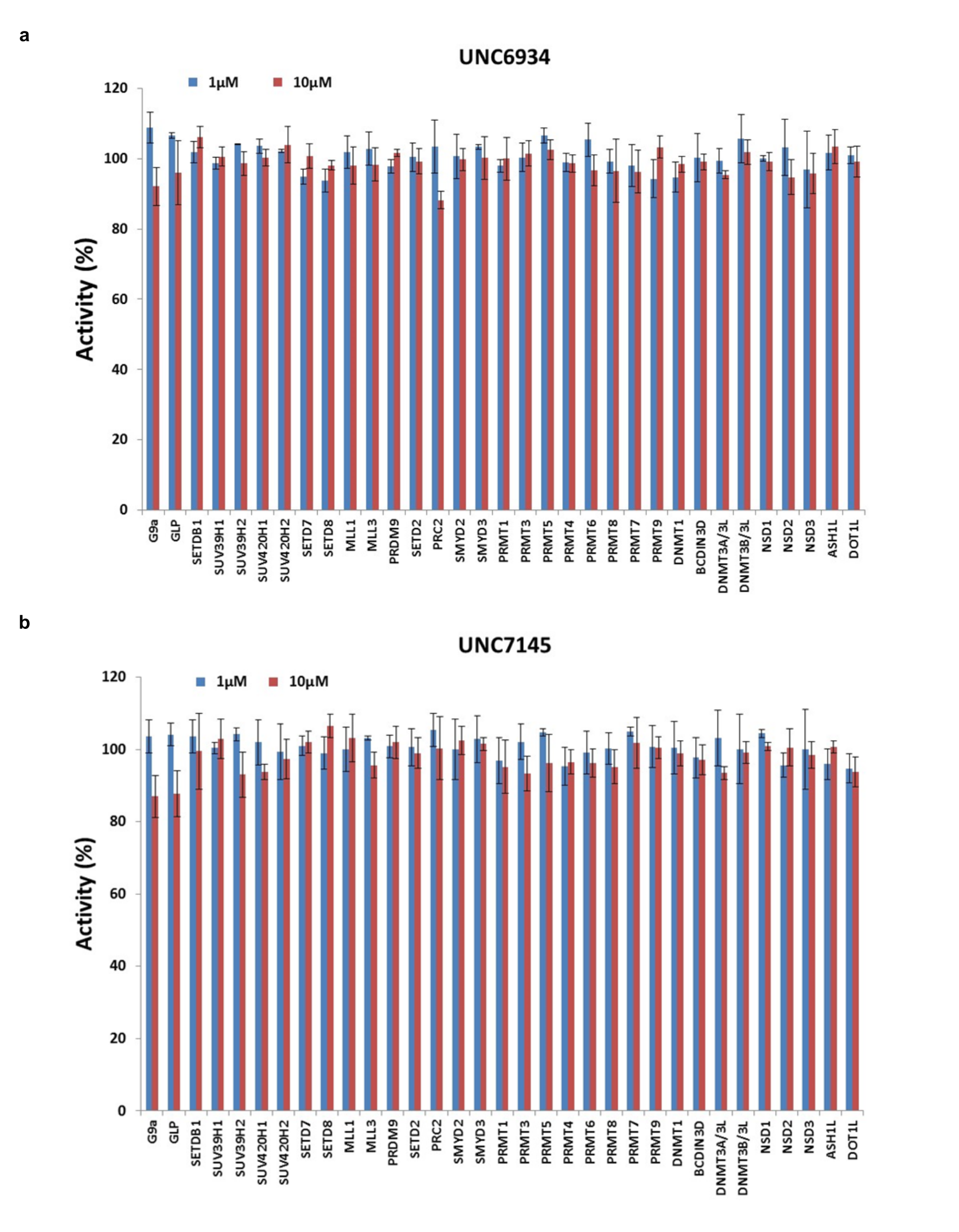

**Supplementary Figure 3**: **Selectivity profile.** Inhibitory activity 1 and 10 µM (blue and red resp.) of UNC6934 (**a**) and UNC7145 (**b**) on a panel of 33 protein methyltransferases. Constructs used for NSD1, NSD2 and NSD3 correspond to the catalytic domain (residues 1810-2120, 934-1241 and 1014-1323 respectively).

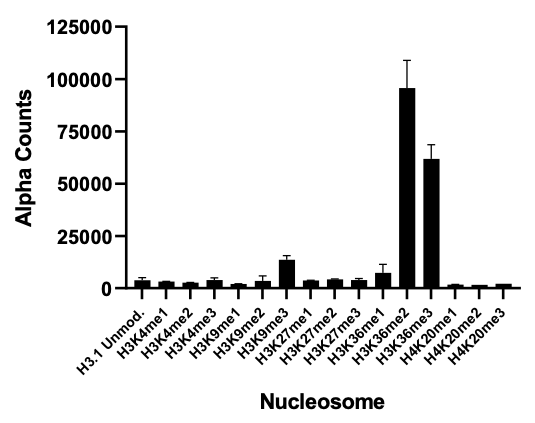

**Supplementary Figure 4: Nucleosome binding data obtained by AlphaScreen.** (**a**) Binding profile of NSD2 PWWP1 against a panel of methylated designer nucleosomes.

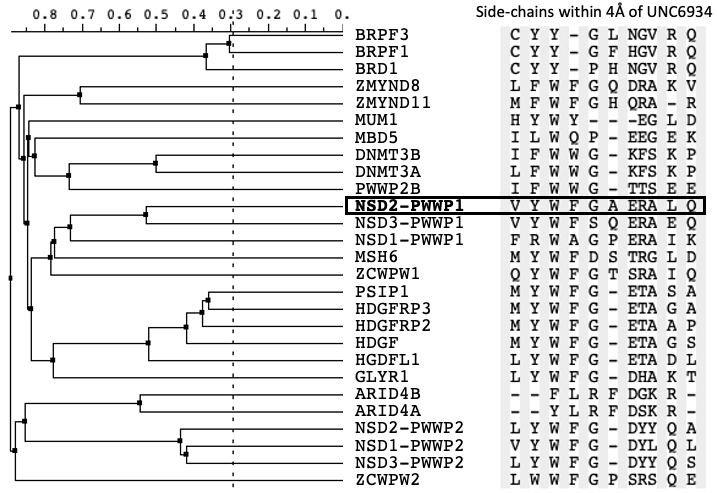

**Supplementary Figure 5: Structural diversity of the UNC6934 binding pocket.** Sequence alignment dendogram of human PWWP domains focused on residues within 4Å of **UNC6934** (generated with ICM (Molsoft, San Diego)).

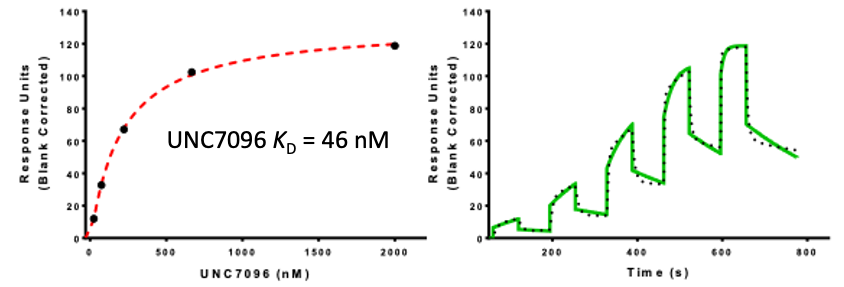

**Supplementary Figure 6: Binding of UNC7096 to NSD2-PWWP1.** SPR sensorgram (solid green) is shown with the kinetic fit (black dots), and K_d_ values were generated from kinetic fitting; The steady state responses (black circles) were shown with the steady state 1:1 binding model fitting (red dashed line). NSD2-PWWP1 domain was immobilized on the flow cell of an CM5 sensor chip in 1x HBS-EP buffer, yielding ̴4000 RU. Using buffer with 0.5% DMSO and single cycle kinetic with 60 s contact time and a dissociation time of 120s at a flow rate of 75 µL/min.

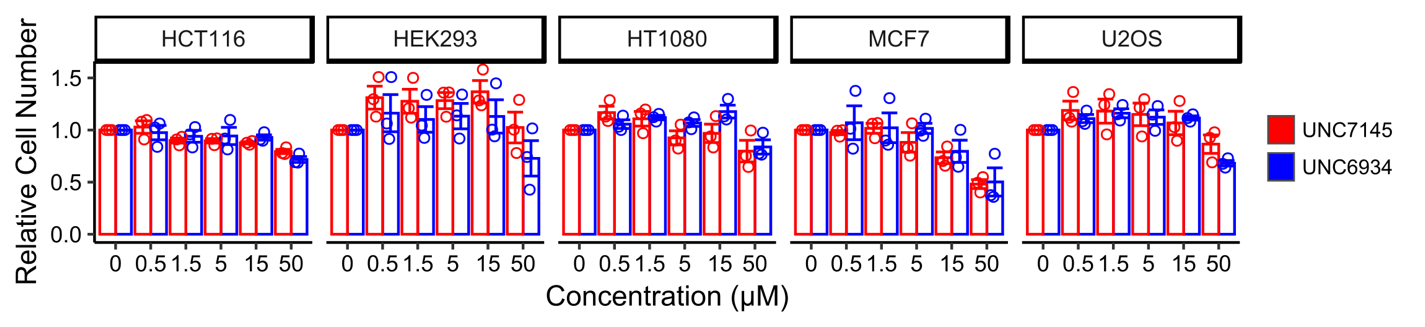

**Supplementary Figure 7: Cytotoxicity Profiling of UNC6934 and UNC7145.** Effect of **UNC6934** and **UNC7145** on cell viability after a 72-hour treatment. Cells were treated with indicated concentrations of either **UNC7145** or **UNC6934** in 96-well plates and nuclei counted by staining Vybrant™ DyeCycle™ Green Stain and imaging on an IncuCyte live-cell analysis system. Each point represents the average number of cells across for fields relative to DMSO treated control (n =3).

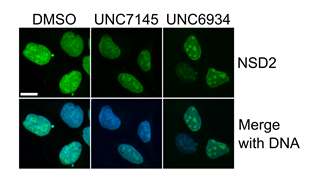

**Supplementary Figure 8: UNC6934 promotes nucleolar localization of NSD2.** Confocal microscopy of U2OS cells treated for 4 hours with 5 µM **UNC6934** or **UNC7145** stained for NSD2 (green) and DNA (blue; Hoescht 33342), scale bar = 15 µm.

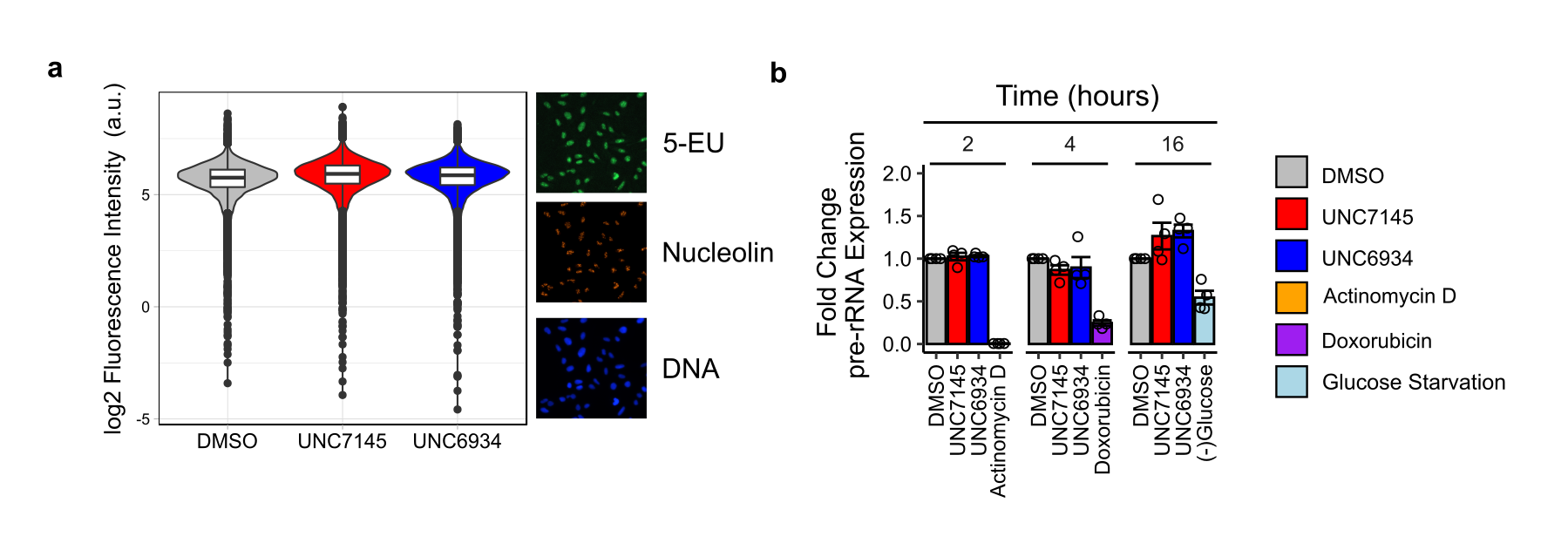

**Supplementary Figure 9: UNC6934 has no significant effect on pre-rRNA expression.** (**a**) 5-EU incorporation assay to measure nascent rRNA synthesis in cells pre-treated for 1 hour with 5 µM **UNC6934** or **UNC7145**. On the left, violin and boxplots generated from three independent experiments show the log2 fluorescence intensity of 5-EU signal within nucleolar regions defined by staining with a nucleolin antibody per nuclei. On the right, representative microscopy images captured on an EVOS FL Auto 2 microscope are shown. (**b**) pre-rRNA expression levels measured by RT-qPCR show no significant effects on ribosome transcription in response to 5 µM **UNC6934** or **UNC7145** at 2, 4, or 16 hours. Expression levels of pre-rRNA shown are relative to DMSO control and normalized to the beta-2-microglobulin housekeeping gene. Each time point represents an independent experiment with four technical replicates. Included in each time point is a control treatment known to disrupt rRNA expression^43,47,48^, controls include at 2 hours - 250 nM actinomycin D, at 4 hours - 1 µM doxorubicin, and at 16 hours - glucose starvation. Primer sequences are shown in Supplementary Table 3

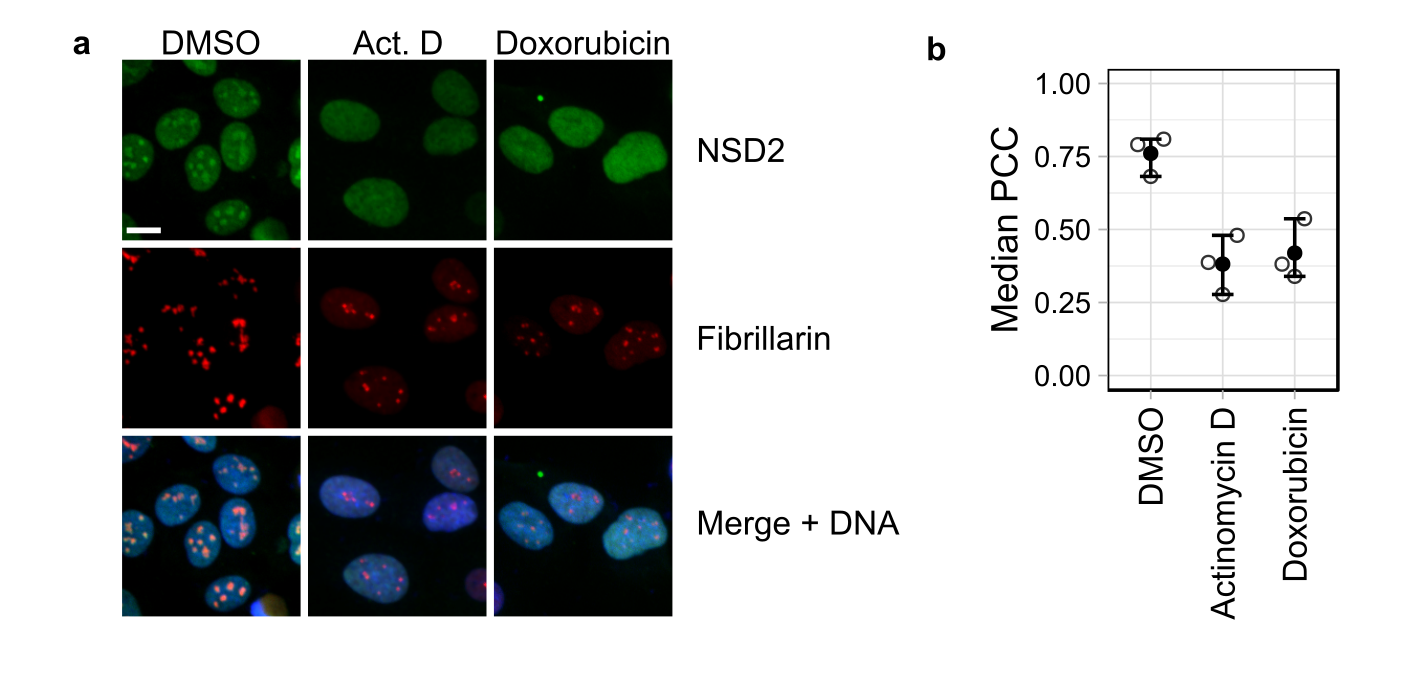

**Supplementary Figure 10: Nucleolar perturbation alters the localization of NSD2.** (**a**) Representative confocal microscopy images of NSD2 and fibrillarin staining in U2OS cells treated for four hours with DMSO control, 50 nM actinomycin D, or 1 µM doxorubicin (scale bar = 15 µm). (**b**) Quantification of co-localization between NSD2 and the nucleolar marker fibrillarin as determined by Pearson correlation coefficient (PCC). The median Pearson Correlation Coefficient is shown for three independent biological replicates (n = 3).

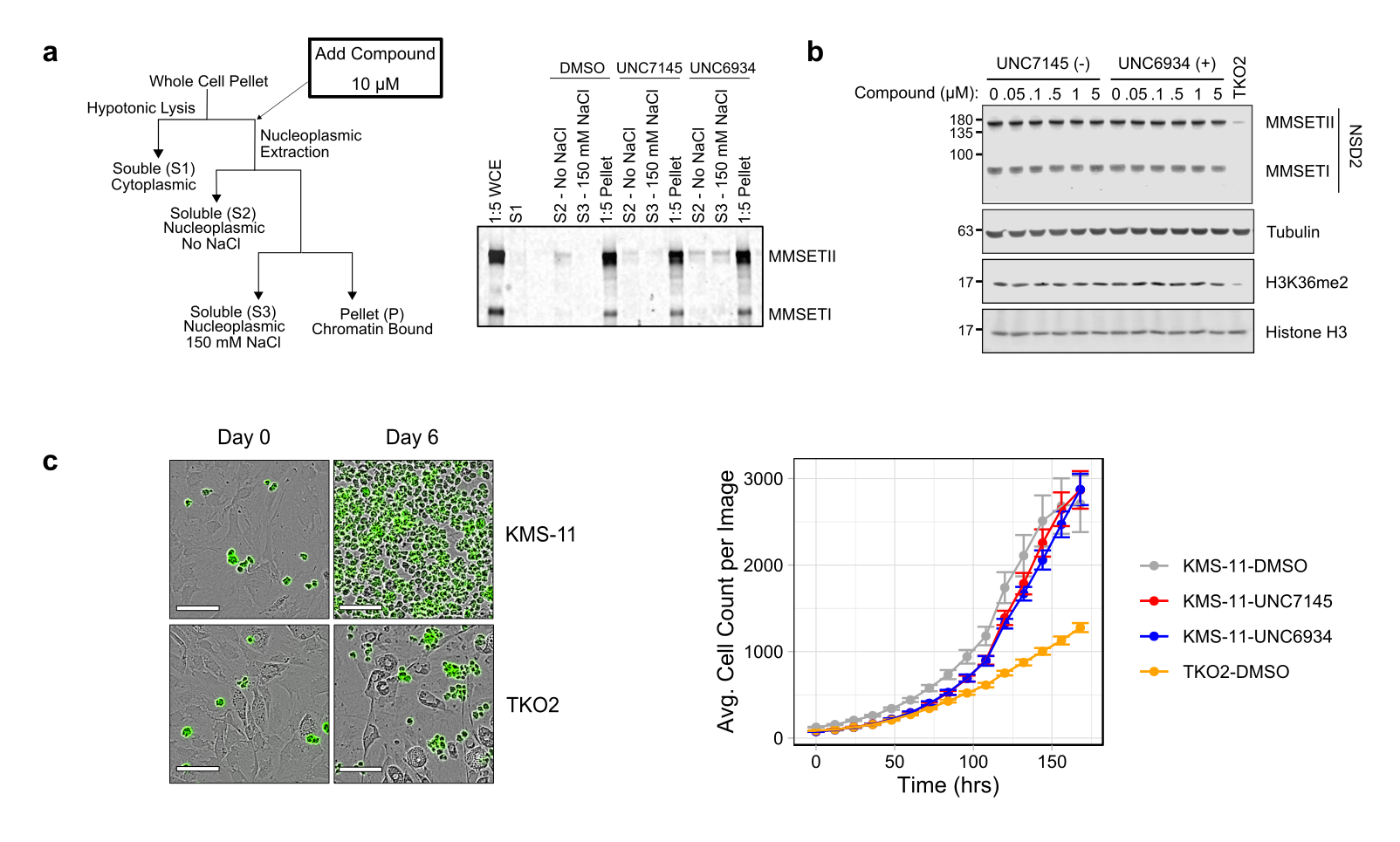

**Supplementary Figure 11: Effect of UNC6934 on KMS11 t(4;14) multiple myeloma cell line and global H3K36me2 levels.** (**a**) Cellular fractionation experiments in KMS-11 cells show increased displacement of NSD2 from chromatin in the S3 fraction containing 150 mM NaCl relative to DMSO or **UNC7145** treated cell lysates. On the left, a schematic of the fractionation protocol. On the right, western blot analysis of NSD2 in each fraction. 1:5 indicates a fifth of the sample was run on the SDS-PAGE relative to other samples. (**b**) Western blot analysis of global H3K36me2 levels in KSM11 cells treated for 72 hours with several doses of either **UNC7145** or **UNC6934** shows no significant change in response to compound. A well-characterized isogenic line harboring a deletion of exon 7 in the mutated KMS11 NSD2 allele is included as a control. (**c**) **UNC6934** does not affect the proliferation of KMS11 cells on bone marrow stroma *in vitro*. Stable GFP expressing KMS11 and isogenic TKO lines were pre-treated with 5 µM of either **UNC7145** or **UNC6934** for 10 days prior to plating on a confluent layer of the OP9 murine bone stromal cell line. Proliferation was monitored by measuring GFP^+^ cells over the course of seven days on an IncuCyte live cell analysis system. Representative images are shown on left. On the right, the average cell count per image is shown for two independent experiments, each with at least six technical replicates.

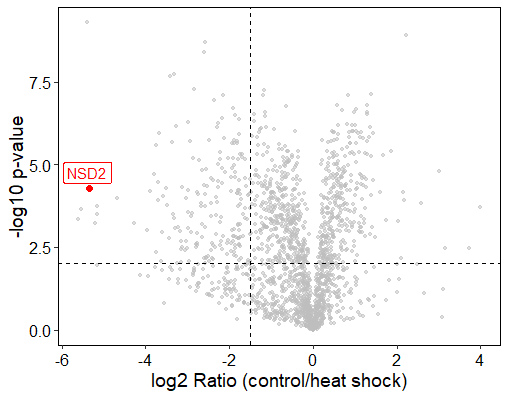

**Supplementary Figure 12. NSD2 Accumulates in the Nucleolus in Response to Heat Shock.** Volcano plot of published data^31^ shows nucleolar proteins in untreated and heat-shocked K562 cells with NSD2 highlighted in red as significantly enriched. The authors performed statistical analysis using Student’s t-test (false discovery rate (FDR) < 0.01; fold change (FC) > 2).

**Supplementary Table 1.** Crystallographic data and refinement statistics

|  | NSD2-PWWP1+MRT866 | NSD2-PWWP1+**UNC6934** |
| --- | --- | --- |
| **PDB Code** | 7LMT | 6XCG |
| **Data collection** |  |  |
| Space group | P1 | P1 |
| Cell dimensions |  |  |
| *a*, *b*, *c* (Å) | 68.3, 69.1, 80.2 | 49.6, 50.0, 51.2 |
| *α*, *β*, *γ* (°) | 75.5,79.4,60.5 | 91.4,91.9,118.3 |
| Resolution (Å) (highest resolution shell) | 48.81-2.26(2.33-2.26) | 50.00-1.64(1.67-1.64) |
| Unique reflections | 51608 | 49043 |
| *R*_merge_ | 6.2(98.2) | 5.3(43.6) |
| *I*/σ*I* | 8.6(0.9) | 37.9(1.9) |
| Completeness(%) | 89.9(82.0) | 92.7(86.5) |
| Redundancy | 2.1(2.1) | 5.0(2.6) |
| **Refinement** |  |  |
| Resolution (Å) | 43.3-2.27 | 25.56-1.64 |
| No. reflections (test set) | 51137(2502) | 47761(1168) |
| *R*_work/_ *R*_free_ (%) | 21.8/26.0 | 17.8/22.2 |
| No. atoms |  |  |
| Protein | 7478 | 3180 |
| Compound | 184 | 99 |
| Water | 14 | 608 |
| B-factors (Å^2^) |  |  |
| Protein | 71.0 | 22.5 |
| Compound | 61.3 | 24.8 |
| Water | 34.3 | 31.1 |
| RMSD |  |  |
| Bond lengths (Å) | 0.01 | 0.011 |
| Bond angles (º) | 1.06 | 1.434 |
| Ramachandran plot % residues |  |  |
| Favored | 99.0 | 99.5 |
| Additional allowed | 1.0 | 0.5 |
| Generously allowed | 0 | 0.0 |
| Disallowed | 0 | 0.0 |

**Supplementary Table 2.** GPCR Selectivity Data

| **Compound** | **Receptor** | **Inhibition 1** | **Inhibition 2** | **Inhibition 3** | **Inhibition 4** | **Mean %** | **Ki (nM)** |
| --- | --- | --- | --- | --- | --- | --- | --- |
| **UNC6934** | 5-HT1A | 23.89 | 1.72 | -0.07 | 27.68 | 13.31 |  |
| **UNC6934** | 5-HT1B | 10.33 | -2.42 | -0.58 | -3.75 | 0.9 |  |
| **UNC6934** | 5-HT1D | 6.2 | 15.98 | 10.6 | 12.16 | 11.24 |  |
| **UNC6934** | 5-HT1E | 2.09 | 9.97 | -3.74 | -11.49 | -0.79 |  |
| **UNC6934** | 5-HT2A | 11.6 | 22.44 | 13.9 | 42.9 | 22.71 |  |
| **UNC6934** | 5-HT2B | 1.7 | 4.62 | 4.48 | 12.52 | 5.83 |  |
| **UNC6934** | 5-HT2C | 19.25 | 27.09 | 20.15 | 25.53 | 23.01 |  |
| **UNC6934** | 5-HT3 | 8.3 | -48.2 | -23.33 | 1.49 | -15.44 |  |
| **UNC6934** | 5-HT5A | 12.19 | 1.38 | 9.11 | 12.97 | 8.91 |  |
| **UNC6934** | 5-HT6 | 8.82 | 15.47 | -1.64 | -15.59 | 1.77 |  |
| **UNC6934** | 5-HT7A | -9.65 | -16.68 | 1.5 | -2.27 | -6.78 |  |
| **UNC6934** | Alpha1A | 20.43 | 18.61 | -3.86 | 19.06 | 13.56 |  |
| **UNC6934** | Alpha1B | 40.57 | 24.76 | 24.89 | 22.77 | 28.25 |  |
| **UNC6934** | Alpha1D | 4.23 | 13.02 | 3.05 | 57.98 | 19.57 |  |
| **UNC6934** | Alpha2A | 4.73 | -24.1 | -17.27 | 18.38 | -4.57 |  |
| **UNC6934** | Alpha2B | 9.74 | 2.91 | 11.02 | 29.31 | 13.25 |  |
| **UNC6934** | Alpha2C | 19.32 | 4.02 | 4.71 | 30.21 | 14.57 |  |
| **UNC6934** | Beta1 | -3.98 | -3.07 | -7.91 | -20.94 | -8.98 |  |
| **UNC6934** | Beta2 | 13.97 | -5.56 | 82.85 | 6.37 | 24.41 |  |
| **UNC6934** | Beta3 | 30.07 | 24.21 | -15.58 | -2.12 | 9.15 |  |
| **UNC6934** | BZP Rat Brain Site | -4.03 | -3.8 | -17.77 | -12.59 | -9.55 |  |
| **UNC6934** | D1 | 6.69 | 3.77 | -2.6 | 5.23 | 3.27 |  |
| **UNC6934** | D2 | -4.48 | -4.31 | -10.72 | -6.91 | -6.61 |  |
| **UNC6934** | D3 | 11.52 | 11.17 | 20 | 23.53 | 16.56 |  |
| **UNC6934** | D4 | -40.11 | 22.04 | -4.7 | 10.84 | -2.98 |  |
| **UNC6934** | D5 | 1.99 | -18.91 | -17.91 | -11.61 | -11.61 |  |
| **UNC6934** | DAT | 58.31 | 37.73 | 29.93 | 50.18 | 44.04 |  |
| **UNC6934** | DOR | 10.82 | 10.22 | 15.29 | 33.28 | 17.4 |  |
| **UNC6934** | GABAA | 11.16 | 15.2 | 4.38 | 12.26 | 10.75 |  |
| **UNC6934** | H1 | 54.96 | 42.68 | 35.32 | 43.17 | 44.03 |  |
| **UNC6934** | H2 | 35.37 | 22.4 | 31.25 | 0 | 22.26 |  |
| **UNC6934** | H3 | -32.92 | -48.2 | -14.64 | -41.91 | -34.42 | >10,000 |
| **UNC6934** | H4 | 5.69 | 0.91 | 2.24 | -1.73 | 1.78 |  |
| **UNC6934** | KOR | -3.75 | -8.91 | 11.91 | 0.42 | -0.08 |  |
| **UNC6934** | M1 | -10.22 | -17.92 | -17.51 | 9.88 | -8.94 |  |
| **UNC6934** | M2 | 32.73 | -6.4 | -15.42 | 5.35 | 4.07 |  |
| **UNC6934** | M3 | 23.96 | 6.74 | 13.69 | 22.3 | 16.67 |  |
| **UNC6934** | M4 | -32.86 | -18.65 | -29.66 | -0.18 | -20.34 |  |
| **UNC6934** | M5 | 15.73 | -3.16 | -25.62 | -7.44 | -5.12 |  |
| **UNC6934** | MOR | 58.55 | 33.8 | 60.34 | 46.59 | 49.82 |  |
| **UNC6934** | NET | -19.48 | -14.89 | -34.9 | -2.91 | -18.05 |  |
| **UNC6934** | PBR | 50.16 | 55.79 | 119.56 | 133.39 | 89.73 | >10,000 |
| **UNC6934** | SERT | 59.39 | 55.69 | 62.25 | 62.37 | 59.93 | 1394 |
| **UNC6934** | Sigma 1 | 12.15 | 30.62 | 23.33 | 40.83 | 26.73 |  |
| **UNC6934** | Sigma 2 | 10.74 | -11.26 | 12.66 | 2.51 | 3.66 |  |
| **UNC7145** | 5-HT1A | 27.48 | -5.07 | 4.92 | 15.9 | 10.81 |  |
| **UNC7145** | 5-HT1B | 12.5 | 10.83 | -6.83 | 1.67 | 4.54 |  |
| **UNC7145** | 5-HT1D | 8.61 | 5.21 | 22.08 | 5.35 | 10.31 |  |
| **UNC7145** | 5-HT1E | 4.76 | 31.55 | -6.03 | 1.84 | 8.03 |  |
| **UNC7145** | 5-HT2A | 10.51 | -6.85 | 0.52 | -29.4 | -6.31 |  |
| **UNC7145** | 5-HT2B | 5.79 | 8.13 | 3.45 | 6.23 | 5.9 |  |
| **UNC7145** | 5-HT2C | 30.91 | 1.77 | 12.75 | 47.94 | 23.34 |  |
| **UNC7145** | 5-HT3 | -12.2 | -17.82 | -13.61 | 3.54 | -10.02 |  |
| **UNC7145** | 5-HT5A | -4.54 | 17.86 | 0.61 | -0.42 | 3.38 |  |
| **UNC7145** | 5-HT6 | 6.67 | 29.26 | 8.16 | 31.25 | 18.84 |  |
| **UNC7145** | 5-HT7A | -14.02 | 3.04 | -14.28 | -5.79 | -7.76 |  |
| **UNC7145** | Alpha1A | 7.26 | 2.5 | -3.18 | -1.59 | 1.25 |  |
| **UNC7145** | Alpha1B | 6.29 | 19.45 | 14.93 | 28.08 | 17.19 |  |
| **UNC7145** | Alpha1D | 11.32 | 14.34 | 14.94 | 39.15 | 19.94 |  |
| **UNC7145** | Alpha2A | -14.24 | -20.74 | -0.26 | 15.24 | -5 |  |
| **UNC7145** | Alpha2B | 3.03 | -3.23 | 8.47 | 31.28 | 9.89 |  |
| **UNC7145** | Alpha2C | 28.82 | 8.6 | -14.3 | 25.71 | 12.21 |  |
| **UNC7145** | Beta1 | 6.02 | 14.8 | -52.14 | -3.07 | -8.6 |  |
| **UNC7145** | Beta2 | 1.49 | 40.27 | 70.37 | 8.27 | 30.1 |  |
| **UNC7145** | Beta3 | 5.37 | -5.28 | 5.95 | 16.37 | 5.6 |  |
| **UNC7145** | BZP Rat Brain Site | 5.44 | -25.88 | -7.41 | -3.35 | -7.8 |  |
| **UNC7145** | D1 | -2.7 | -1.76 | -6.35 | 0.22 | -2.65 |  |
| **UNC7145** | D2 | -9.68 | -12.97 | -0.67 | -9.68 | -8.25 |  |
| **UNC7145** | D3 | 27.42 | -50.29 | 36.25 | 34.83 | 12.05 |  |
| **UNC7145** | D4 | 1.99 | 8.49 | -5.24 | 18.07 | 5.83 |  |
| **UNC7145** | D5 | 14.26 | -9.29 | 3.65 | -12.6 | -1 |  |
| **UNC7145** | DAT | 1.22 | -15.04 | 4.54 | 6.87 | -0.6 |  |
| **UNC7145** | DOR | 19.03 | 14.81 | 54.54 | 42.46 | 32.71 |  |
| **UNC7145** | GABAA | 3.46 | 10.06 | 13 | 10.8 | 9.33 |  |
| **UNC7145** | H1 | 58.15 | 74.1 | 54.96 | 72.14 | 64.84 | >10,000 |
| **UNC7145** | H2 | 15.1 | -3.62 | 2.94 | 5.85 | 5.07 |  |
| **UNC7145** | H3 | -5.06 | -8.35 | -33.22 | 0 | -11.66 |  |
| **UNC7145** | H4 | 8.13 | 1.73 | 9.35 | 4.88 | 6.02 |  |
| **UNC7145** | KOR | -0.58 | -9.08 | 3.25 | -4.08 | -2.62 |  |
| **UNC7145** | M1 | -10.22 | -9.47 | 7.29 | -8.25 | -5.16 |  |
| **UNC7145** | M2 | 3.1 | -64.69 | -15.74 | 0.85 | -19.12 |  |
| **UNC7145** | M3 | 1.01 | 4.19 | -4.98 | 9.48 | 2.43 |  |
| **UNC7145** | M4 | 45.8 | 9.36 | -24.04 | -34.16 | -0.76 |  |
| **UNC7145** | M5 | 56.73 | 14.3 | -2.81 | -13.5 | 13.68 |  |
| **UNC7145** | MOR | 67.07 | 60.61 | 74.77 | 61.99 | 66.11 | 7950 |
| **UNC7145** | NET | -48.79 | 0 | -30.44 | 4.22 | -18.75 |  |
| **UNC7145** | PBR | 67.57 | 91.65 | 117.51 | 134.92 | 102.91 | >10,000 |
| **UNC7145** | SERT | 61.06 | 20.02 | 41.02 | 37.08 | 39.8 |  |
| **UNC7145** | Sigma 1 | 1.94 | -1.94 | 4.37 | 39.85 | 11.06 |  |
| **UNC7145** | Sigma 2 | 12.51 | -10.41 | -24.56 | 9.2 | -3.32 |  |

**Supplementary Table 3**. Primer Sequences for pre-rRNA Expression Analysis

| **Target** | **Forward** | **Reverse** | **Reference** |
| --- | --- | --- | --- |
| B2M | GAGTGCTGTCTCCATGTTTGATGT | AAGTTGCCAGCCCTCCTAGAG | ^49^ |
| pre-rRNA | GCCTTCTCTAGCGATCTGAGAG | CCATAACGGAGGCAGAGACA | ^50^ |

**Compound purity:**

QC Data for MRT866 Purchased from Enamine (Cat. No. Z357319716)

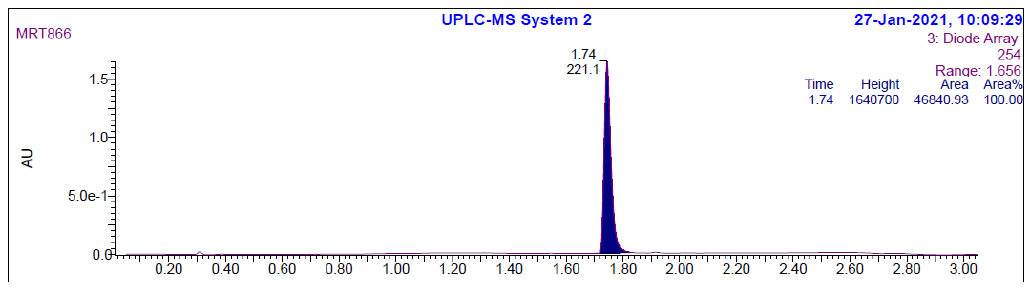

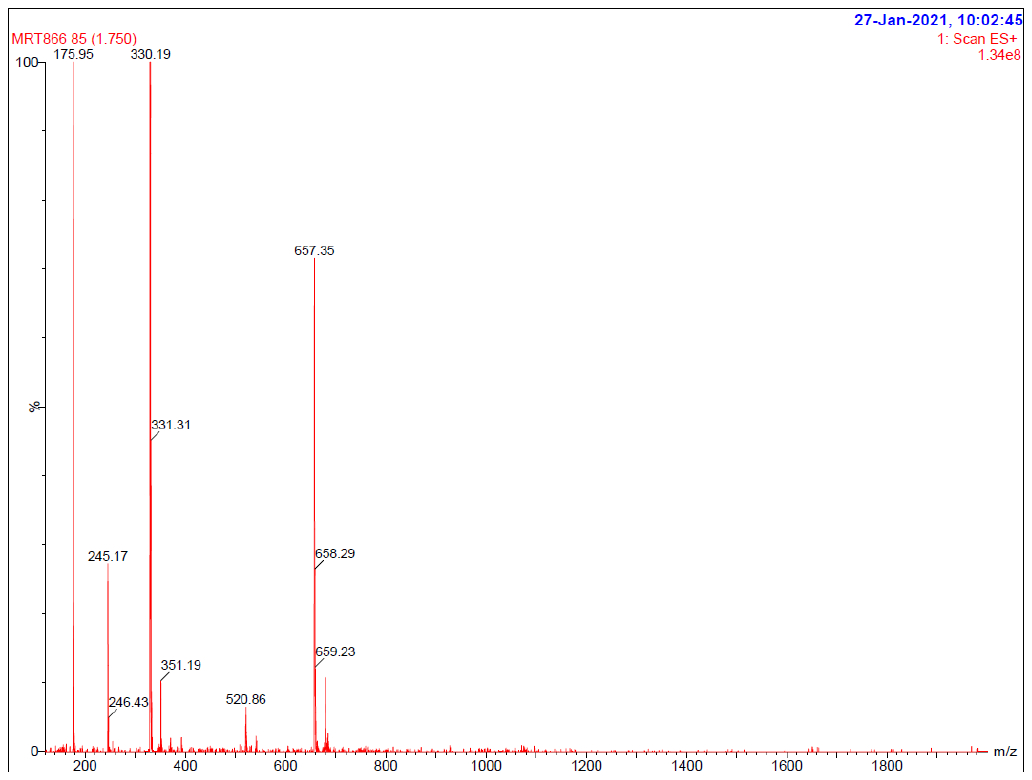

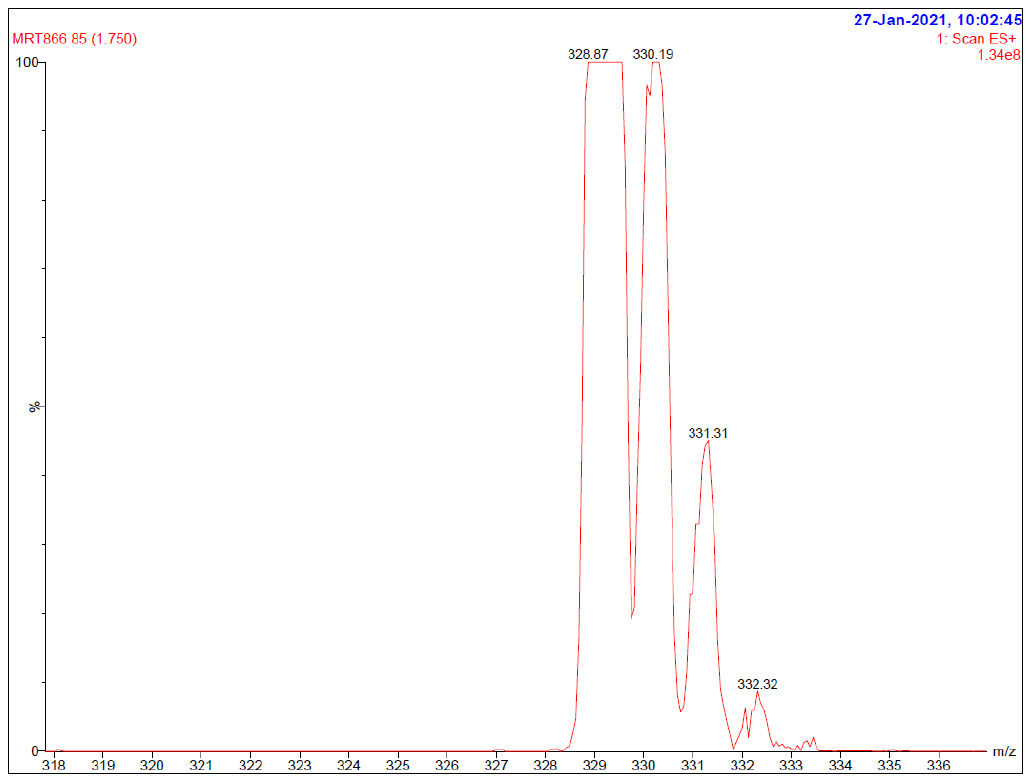

QC Data for UNC6934

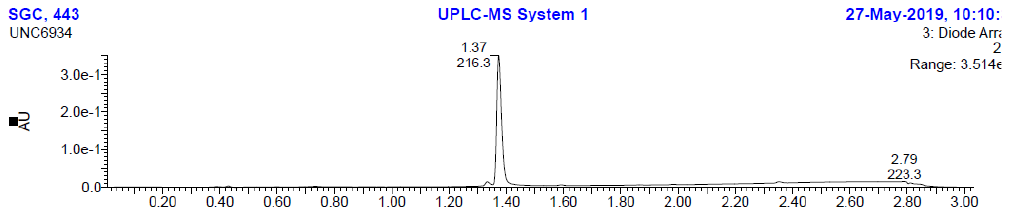

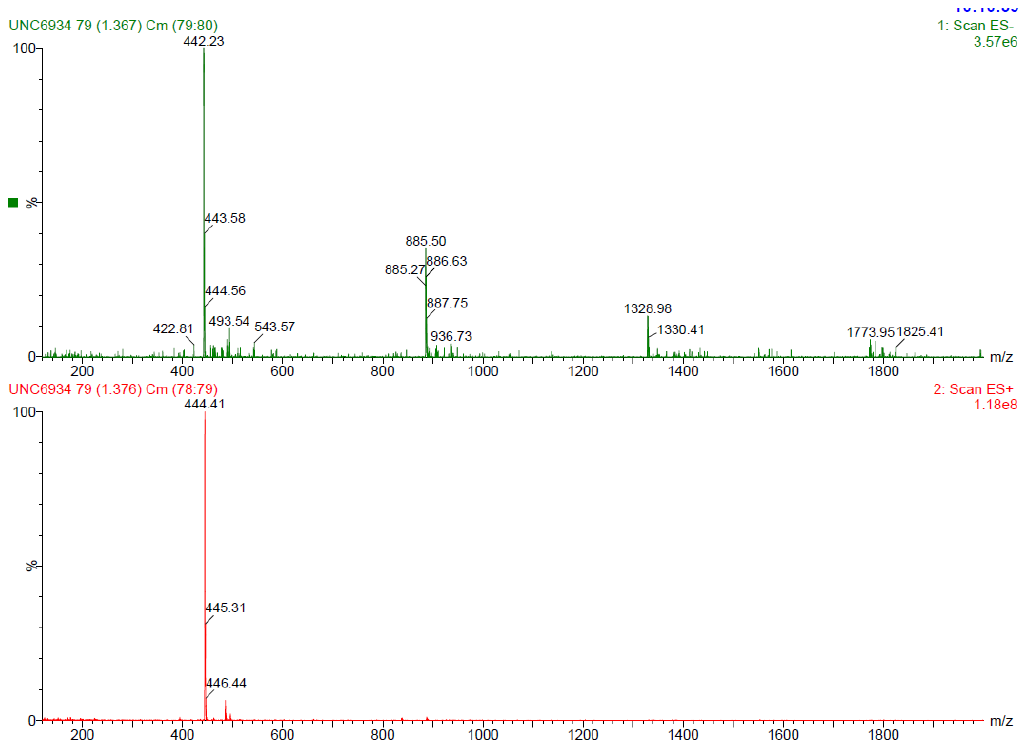

QC Data for UNC7145

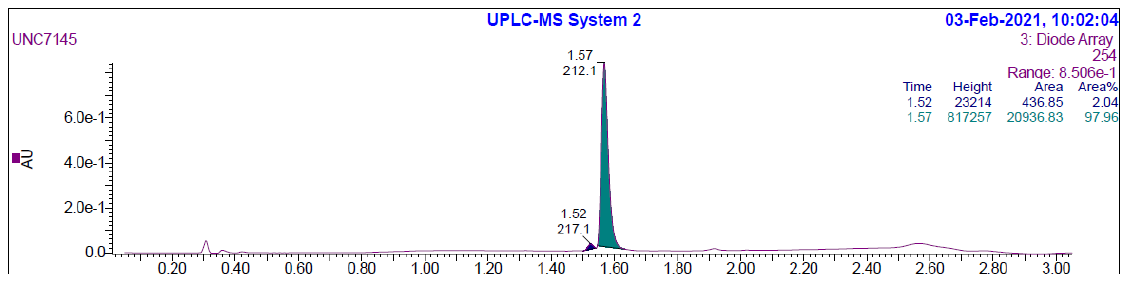

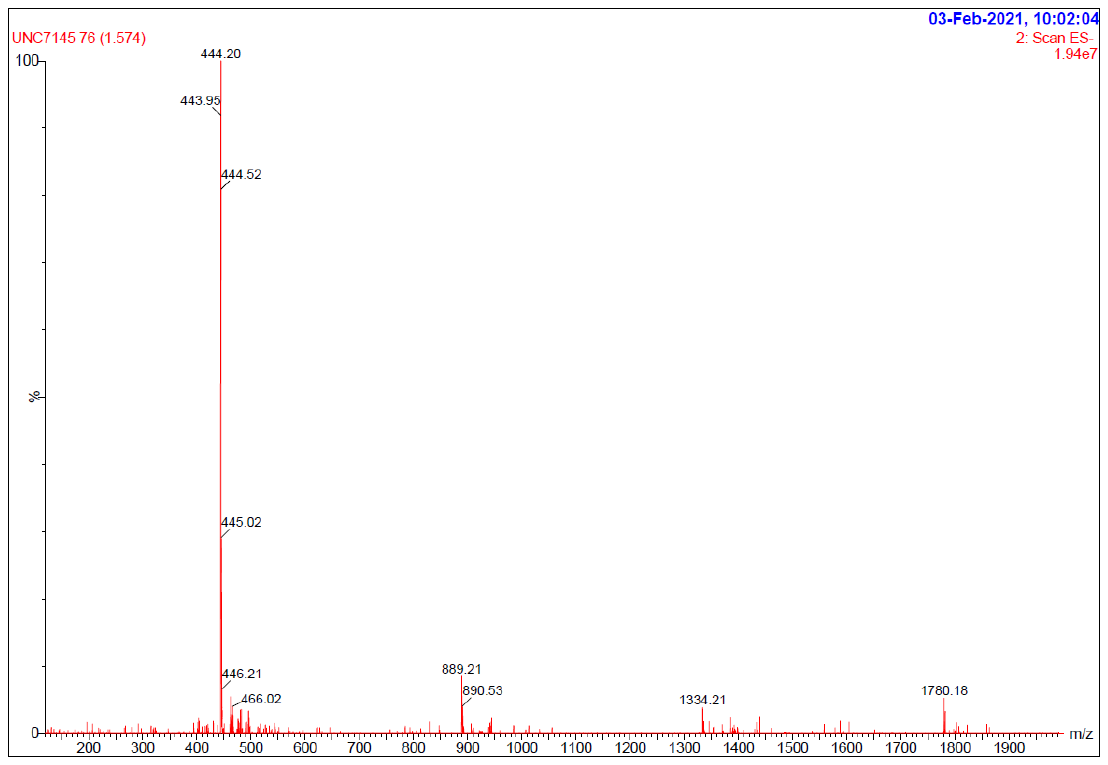

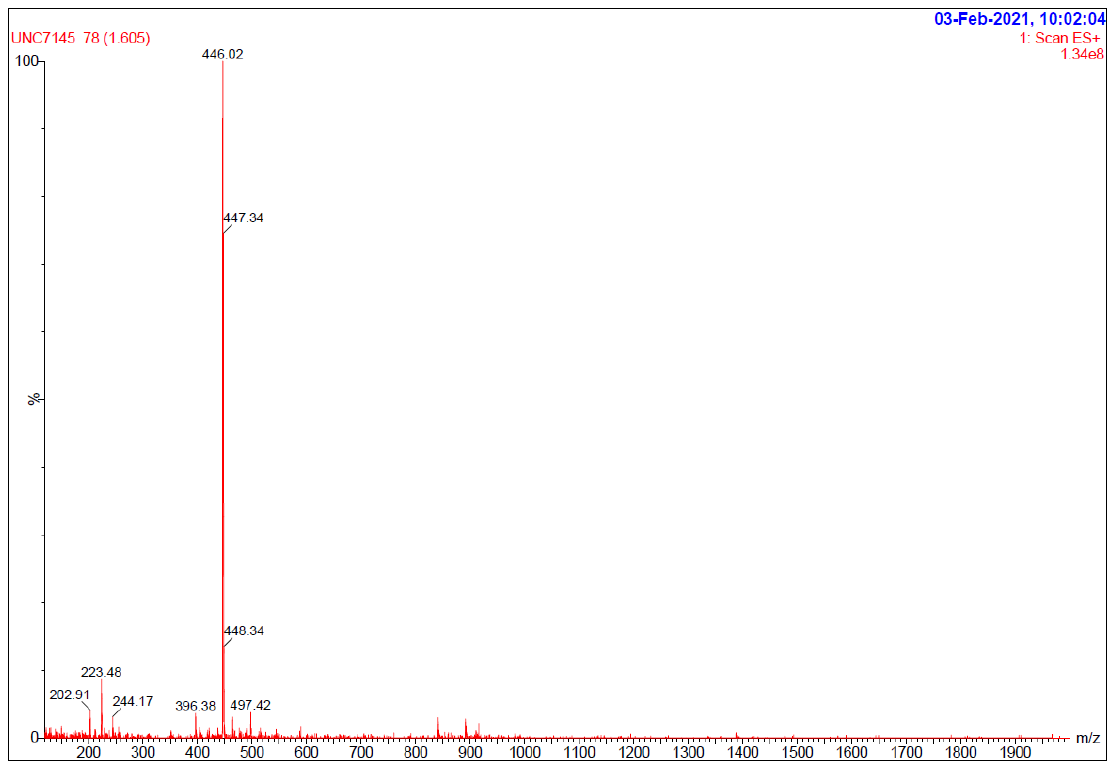
